# The Mut^+^ strain of *Komagataella phaffii* (*Pichia pastoris*) expresses P*_AOX1_* 5 and 10 times faster than Mut^s^ and Mut^−^ strains: Evidence that formaldehyde or/and formate are true inducers of AOX

**DOI:** 10.1101/573519

**Authors:** Anamika Singh, Atul Narang

## Abstract

The methylotrophic yeast *Komagataella phaffii* is among the most popular hosts for recombinant protein synthesis. Most recombinant proteins were expressed in the wild-type Mut^+^ host strain from the methanol-inducible promoter P_*AOX1*_. Since methanol metabolism has undesirable consequences, two additional host strains, Mut^s^ (*AOX1*^*-*^) and Mut^−^ (*AOX1*^*-*^ *AOX2*^*-*^), were introduced which consume less methanol and reportedly also express recombinant protein better than Mut^+^. Both results follow from a simple model based on two widespread assumptions, namely methanol is transported by diffusion and the sole inducer of P_*AOX1*_. To test this model, we studied ^14^C-methanol uptake in the Mut^−^ strain and β-galactosidase expression in all three strains. We confirmed that methanol is transported by diffusion, but in contrast to the literature, Mut^+^ expressed β-galactosidase 5- and 10-fold faster than Mut^s^ and Mut^−^. These results imply that methanol is not the sole inducer of P_*AOX1*_ — metabolites downstream of methanol also induce P_*AOX1*_. We find that formate or/and formaldehyde are probably true inducers since both induce P_*AOX1*_ expression in Mut^−^ which cannot synthesize intracellular methanol from formate or formaldehyde. Formate offers a promising substitute for methanol since it does not appear to suffer from the deficiencies that afflict methanol.

## 1. Introduction

*Komagataella phaffii*, formerly *Pichia pastoris* (Kurtzman, 2005; Mattanovich *et al*., 2009), is an efficient host which has been used for the production of over 500 heterologous proteins (Potvin *et al.*, 2012). It can grow to cell densities of >120 g dry weight (gdw) per litre, it has an efficient secretory system, it is easy to manipulate genetically, and it can perform certain post-translational modifications of heterologous proteins like O-/N-glycosylation and formation of disulphide bonds. However, the most remarkable advantage of this yeast is the presence of a strong, yet tightly controlled, promoter which drives the expression of alcohol oxidase (AOX, EC 1.1.3.13), an enzyme that catalyses the conversion of methanol to formaldehyde (Higgins and Cregg, 1998).

Wild-type *K. phaffii* has two alcohol oxidase genes, *AOX1* and *AOX2*, which operate under the control of their respective promoters, P_*AOX1*_ and P_*AOX2*_. Both promoters display similar regulatory behaviour, but in most expression vectors, P_*AOX1*_ is used to drive the expression of heterologous proteins because it is almost 10 times stronger than P_*AOX2*_ (Cregg *et al.*, 1989).

Although AOX expression is strongly induced by methanol (Cregg *et al.*, 2000), it is desirable to minimize the use of methanol because it is highly flammable. Moreover, its metabolism results in high oxygen demand (Arnau *et al*., 2011), excessive heat generation (Krainer *et al.*, 2012), and formation of toxic products like hydrogen peroxide and formaldehyde (Jungo *et al.*, 2007b).

An elegant solution to these problems was the creation of strains that use methanol sparingly because it serves mainly or exclusively as an inducer of heterologous gene expression rather than as a carbon source for growth. This was achieved by abolishing *AOX1* (resp., *AOX1* and *AOX2*), thus impairing (resp., abrogating) the capacity of cells to consume methanol (Table 1). Cells with the wild-type or “methanol utilization plus” phenotype, denoted Mut^+^, have functional copies of both *AOX1* and *AOX2*. In Mut^s^ or “methanol utilization slow” strains, *AOX1* is disrupted, but *AOX2* is functional, which results in drastically reduced methanol consumption rates. Mut^−^ or “methanol utilization minus” strains express neither *AOX1* nor *AOX2*, and hence cannot consume any methanol. It is therefore necessary to provide Mut^−^ strains with an additional *secondary* carbon source to enable growth (Chiruvolu *et al.*, 1997). It is not necessary, but desirable, to provide Mut^s^ and Mut^+^ strains with a secondary carbon source because its consumption enhances the biomass concentration, and hence volumetric productivity, of the Mut^s^ strain (Brierley *et al*., 1990), and reduces the oxygen demand, heat generation, and toxic product formation of the Mut^+^ strain (Jungo *et al.*, 2007a and Jungo *et al.*, 2007c).

**Table 1:** Phenotype and genotype of three Mut strains (Cregg *et al*., 1987; Tschopp *et al*., 1987b).

| Strain | Genotype | AOX1 | AOX2 | Phenotype |
| --- | --- | --- | --- | --- |
| GS115 | *his4* | Intact | Intact | Mut<sup>+</sup> His<sup>-</sup> |
| KM71 | *aox1Δ::SARG4his4arg4* | Disrupted | Intact | Mut<sup>s</sup> His<sup>-</sup> |
| MC100-3 | *aox1Δ::SARG4aox2Δ::Phis4*<br>*his4arg4* | Disrupted | Disrupted | Mut<sup>-</sup> His<sup>-</sup> |

The Mut^s^ and Mut^−^ strains have drastically lower methanol consumption rates, but it is relevant to ask if this has been achieved at the cost of reduced recombinant protein expression. We are not aware of any study in which the recombinant protein expression of all three strains were compared. However, there are pairwise comparisons of the Mut^+^ strain with the Mut^−^ and Mut^s^ strains, which indicate that even with respect to recombinant protein expression, the Mut^−^ and Mut^s^ strains are superior to the Mut^+^ strain. Chiruvolu *et al.* compared the potential of strains Mut^−^ and Mut^+^ to produce recombinant β-galactosidase protein (Chiruvolu *et al.*, 1997). They found that the Mut^−^ strain provided significantly higher protein yields than the Mut^+^ strain. Since Mut^−^ strains have the added benefit of not consuming methanol, one would expect widespread adoption of these strains, but the potential of this strain has never been explored further. In contrast to the Mut^−^ strain, the Mut^s^ strain has been compared with the Mut^+^ strain in numerous studies which are summarized in Table 2. Although the results of these studies are not unanimous, seven out of ten studies reported better performance of the Mut^s^ strain.

**Table 2:** Comparison of Mut^+^ and Mut^s^ phenotypes as heterologous expression systems.

| Reference | Recombinant protein | Secreted/non-secreted protein | Better strain |
| --- | --- | --- | --- |
| Cregg *et al.*, 1987 | Hepatitis B surface antigen | Non-secreted | Mut<sup>s</sup> |
| Tschopp *et al.*, 1987b | Invertase | Secreted | Mut<sup>s</sup> |
| Brierley *et al.*, 1990;<br>Digan *et al.*, 1989 | Bovine lysozyme | Secreted | Mut<sup>+</sup> |
| Clare *et al.*, 1991 | Tetanus toxin fragment C | Non-secreted | Mut<sup>s</sup> strain containing multiple integrations of the recombinant gene |
| Peng *et al.*, 2004 | Bovine enterokinase light chain | Secreted | Mut<sup>s</sup> |
| Cos *et al.*, 2005 | *Rhizopus oryzae* lipase | Secreted | Mut<sup>s</sup> |
| Pla *et al.*, 2006 | Antibody fragment | Secreted | Mut<sup>s</sup> |
| Kim *et al.*, 2009 | *Coprinus cinerius* peroxidase | Secreted | Mut<sup>+</sup> |
| Krainer *et al.*, 2012 | Horseradish peroxidase C1A | Secreted | Mut<sup>s</sup> |
| Pedro *et al.*, 2015 | Catechol-O-methyltransferase | Non-secreted | Mut<sup>+</sup> |

Interestingly, the superior recombinant protein expression of the Mut^−^ and Mut^s^ strains also follow from the assumptions implicitly made about methanol transport and *AOX1* induction. Specifically, it is assumed that:

1. Methanol enters the cell by diffusion rather than active transport. This assumption, often made implicitly (James Cregg, personal communication), is probably motivated by the observation that in *Saccharomyces cerevisiae*, ethanol enters the cell by diffusion (Loureiro and Ferreira, 1983; Guijarro and Lagunas, 1984), which suggests that methanol, a similar and even smaller compound, also enters *K. phaffii* cells by diffusion.
2. Methanol is the sole inducer of P_*AOX1*_ expression. This assumption is also implicitly used in the literature since methanol has been used to induce P_*AOX1*_ expression in virtually all studies. It probably stems from early studies which explicitly identified methanol as the sole inducer in the methylotrophic yeast *Hansenula polymorpha* (Eggeling and Sahm, 1980).

Now the first assumption implies that if the Mut^−^, Mut^s^, and Mut^+^ strains are exposed to the same concentration of methanol, the steady state intracellular methanol concentrations will be highest in the Mut^−^ strain, intermediate in the Mut^s^ strain, and lowest in the Mut^+^ strain (Fig. 1). Indeed, since the Mut^−^ strain has no AOX, it cannot metabolize methanol, and the steady state concentration of intracellular methanol will be equal to the concentration of extracellular methanol. In the Mut^s^ strain, which possesses low, but significant, AOX levels, the steady state intracellular methanol concentration will be lower than observed in the Mut^−^ strain. Finally, the still higher AOX levels of the Mut^+^ strain will ensure that the steady state intracellular methanol concentration is even lower than that observed in the Mut^s^ strain. Then it follows from the second assumption that the steady state specific expression rate of P_*AOX1*_ is highest in the Mut^−^ strain, slightly lower in Mut^s^ strain, and lowest in the Mut^+^ strain. Thus, in so far as the recombinant protein expression rate is concerned, the Mut^−^ and Mut^s^ strains are superior to the Mut^+^ strain.

**Fig. 1:**
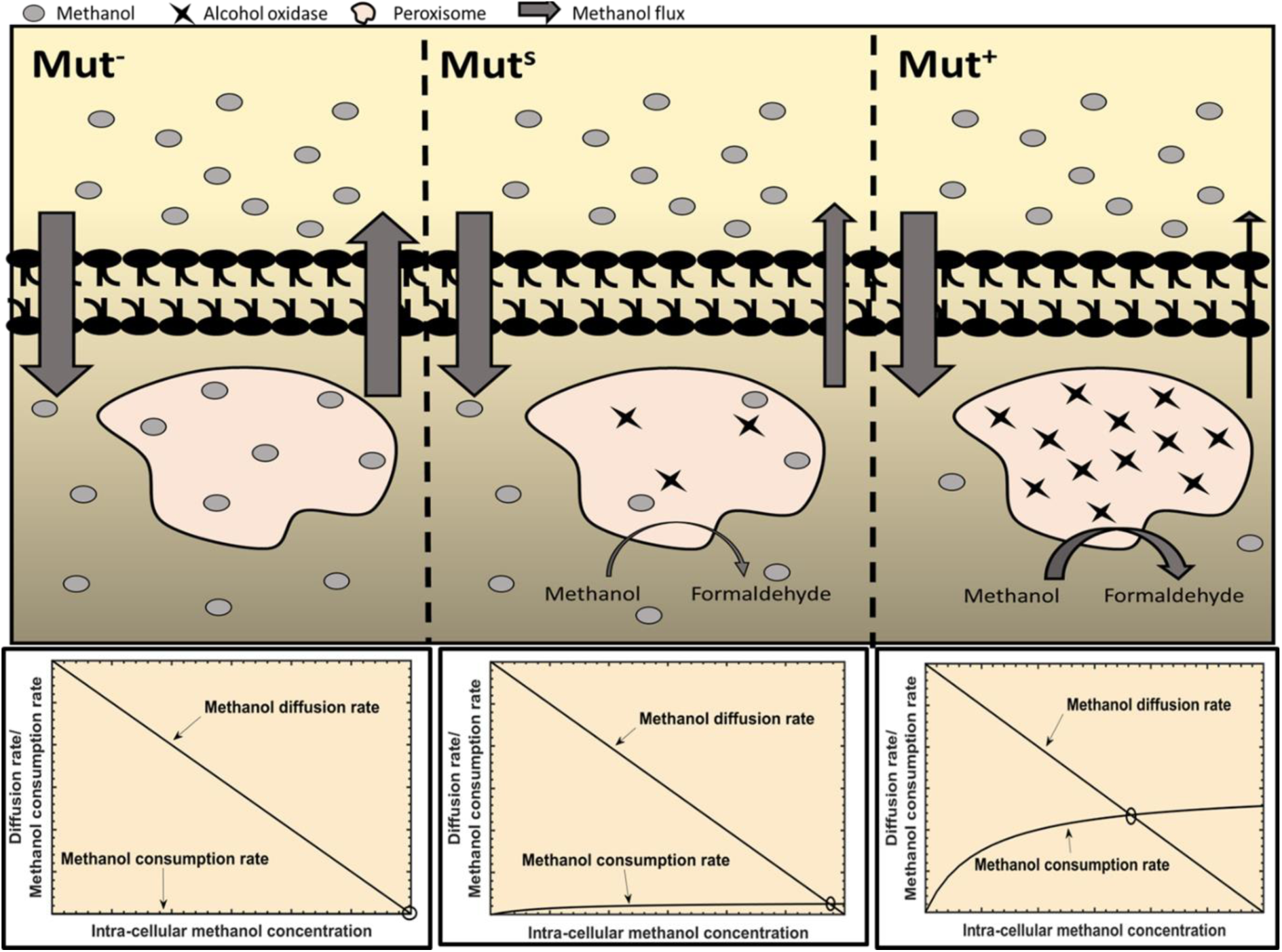
The Mut^−^ strain has the highest steady state intracellular methanol concentration: When Mut^−^, Mut^s^ and Mut^+^ cells are exposed to the same concentration of methanol, the steady state specific activity of AOX, and hence the specific methanol consumption rate, is zero in Mut^−^, small in Mut^s^ and large in Mut^+^. If methanol is transported by diffusion, it follows that the steady state intracellular methanol concentration will be highest in Mut^−^, lower in Mut^s^ and lowest in Mut^+^ strain.

It is useful to test the validity of the above model since, to our knowledge, there has been no attempt to explain the mechanism underlying the observed differences in the expression of the Mut^−^, Mut^s^, and Mut^+^ strains. To this end, we chose β-galactosidase as the recombinant protein because it is easy to assay its specific activity accurately. Moreover, it turns out that since β-galactosidase is neither secreted nor degraded, the specific β-galactosidase expression rates of all three strains, the key variable of our model, can be easily calculated from the corresponding specific β-galactosidase activities. To our surprise, we found that the specific β-galactosidase expression rates observed in the Mut^+^ strain were respectively 5 and 10 times those observed in the Mut^s^ and Mut^−^ strains, thus implying that at least one of the two assumptions of the proposed model is untenable. We show that methanol is transported by diffusion, but not the sole inducer of *AOX1* expression — formaldehyde and formate are also potent inducers of *AOX1* expression. Although formaldehyde inhibits growth strongly, formate is a viable alternative to methanol since it induces P_*AOX1*_ as well as methanol, but does not suffer from the deficiencies that afflict methanol.

## 2. Results

### 2.1 Relationship between specific activity and expression rate of β-galactosidase

In general, a newly synthesized protein is either accumulated, degraded and diluted (by growth) within the cells, or it is secreted out of the cells (Pfeffer *et al*., 2011). More precisely, if *μ*denotes the specific growth rate of the cells (h^−1^), and *p*_*i*_denotes the specific activity of intracellular protein (units gdw^−1^), then the rate of protein accumulation within the cells is given by the mass balance equation

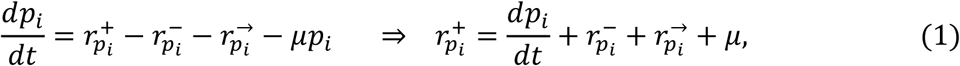

where 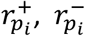 and 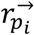 denote the specific rates of protein expression, degradation and secretion, respectively, and *μp*_*i*_denotes the protein dilution rate due to growth. The secreted protein accumulates and degrades in the medium. If *p*_*e*_denotes the concentration of extracellular protein (units L^−1^), then its rate of accumulation is given by the mass balance equation

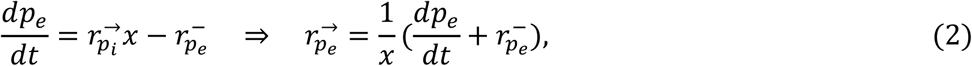

where *x*is the cell density (gdw L^−1^) and 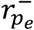 is the degradation rate of extracellular protein (units L^−1^ h^−1^).

Despite the complexity of the above equations, we could easily determine the specific β-galactosidase expression rate, 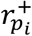 in our experiments. Indeed:

1. The specific secretion rate of β-galactosidase 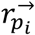 was negligible since we could not detect any β-galactosidase activity in the medium.
2. The specific degradation rate of β-galactosidase 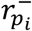 was also negligible. To see this, observe that when fully induced cells are transferred to inducer-free medium, the specific expression rate drops precipitously to its basal level 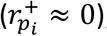, and the subsequent decline of the specific activity due to dilution and degradation is approximated by the equation 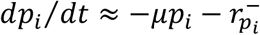. We found that when fully induced Mut^+^ or Mut^−^ cells were transferred to inducer-free medium, the specific β-galactosidase activity declined exponentially, but the decline was entirely due to dilution, i.e, 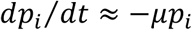 (Fig. S1). It follows that protein degradation is negligible in the absence of the inducer and it can be completely neglected under the reasonable assumption that it does not increase appreciably in the presence of the inducer.
3. We measured the specific β-galactosidase activity in cells that had attained balanced growth by growing exponentially for ∼10 generations. Under these conditions, the specific β-galactosidase activity reaches steady state (*dp*_*i*_/*dt* ≈ 0) and Eq. (2) reduces to the simple form

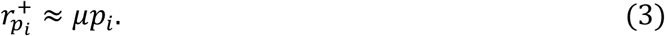

Thus, in our experiments, the specific β-galactosidase expression rate equals the dilution rate, and can be easily calculated from the steady state specific growth rate *μ*and the specific β-galactosidase activity *p*_*i*_.

### 2.2 The specific expression rate of Mut^+^ is 5 and 10 times that of Mut^s^ and Mut^−^

Our goal is to compare the steady state specific β-galactosidase expression rates of the Mut^−^, Mut^s^ and Mut^+^ strains when they are fully induced in the presence of methanol. However, since the Mut^−^ strain cannot grow on methanol, these strains cannot be compared unless the medium contains a secondary carbon source. Furthermore, it is desirable that the mixture of the secondary carbon source and methanol supports a sufficiently high specific growth rate to ensure that the intracellular protein level reaches steady state within a reasonable time, as well as a sufficiently high specific β-galactosidase expression rate to facilitate precise measurement of the specific activity. It turns out that single carbon sources do not satisfy both requirements. Carbon sources supporting high specific growth rates, such as glucose, glycerol and ethanol, strongly repress P_*AOX1*_ even in the presence of methanol, and carbon sources permitting high P_*AOX1*_ expression rates, such as sorbitol, alanine, mannitol and trehalose, support very low specific growth rates in minimal media (Cregg *et al*., 2000; Inan and Meagher, 2001). We found that casamino acids satisfied both requirements, but it did not yield reproducible specific growth rates (data not shown). Supplementation of sorbitol containing minimal medium with yeast nitrogen base (YNB) did not improve the low specific growth rate of 0.03 h^−1^. We were therefore led to consider supplementation of sorbitol containing minimal medium with additional carbon sources. We finally obtained sufficiently high and reproducible specific growth and β-galactosidase expression rates with a mixture of sorbitol (55 mM) and alanine (112 mM), which was used as the secondary carbon source in our experiments.

To determine the steady state expression rates of the Mut^−^, Mut^s^, and Mut^+^ strains when they are fully induced in the presence of methanol, we measured the specific growth rates and β-galactosidase activities of all three strains after ∼10 generations of exponential growth on mixtures of sorbitol, alanine, and various concentrations of methanol. In all three strains, the specific growth rates were essentially independent of the methanol concentration (Fig. S2a) but the specific β-galactosidase activities increased monotonically and plateaued at high methanol concentrations (Fig. S3a). Consequently, the specific β-galactosidase expression rates calculated from Eq. (3) increased hyperbolically and could be fitted to the expression

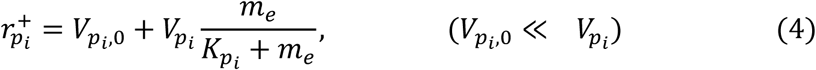

where *m*_*e*_denotes the extracellular methanol concentration; and 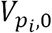 and 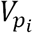 denote the respective specific expression rates under *uninducing* conditions wherein cells grew on sorbitol + alanine, but not methanol (*m*_*e*_= 0), and *fully inducing* conditions wherein cells grew on sorbitol + alanine and saturating concentrations of methanol (*m*_*e*_» *k*_*p*_*i*). Figure 2 shows the specific β-galactosidase specific expression rates of the recombinant Mut^−^, Mut^s^, and Mut^+^ strains under uninducing conditions and fully inducing conditions. Under fully inducing conditions, the specific expression rate of strain Mut^+^ (pSAOH5-T1) was approximately 5 and 10-fold higher than that of strains Mut^s^ (pSAOH5-T1) and Mut^−^ (pSAOH5-T1) respectively.

**Fig. 2:**
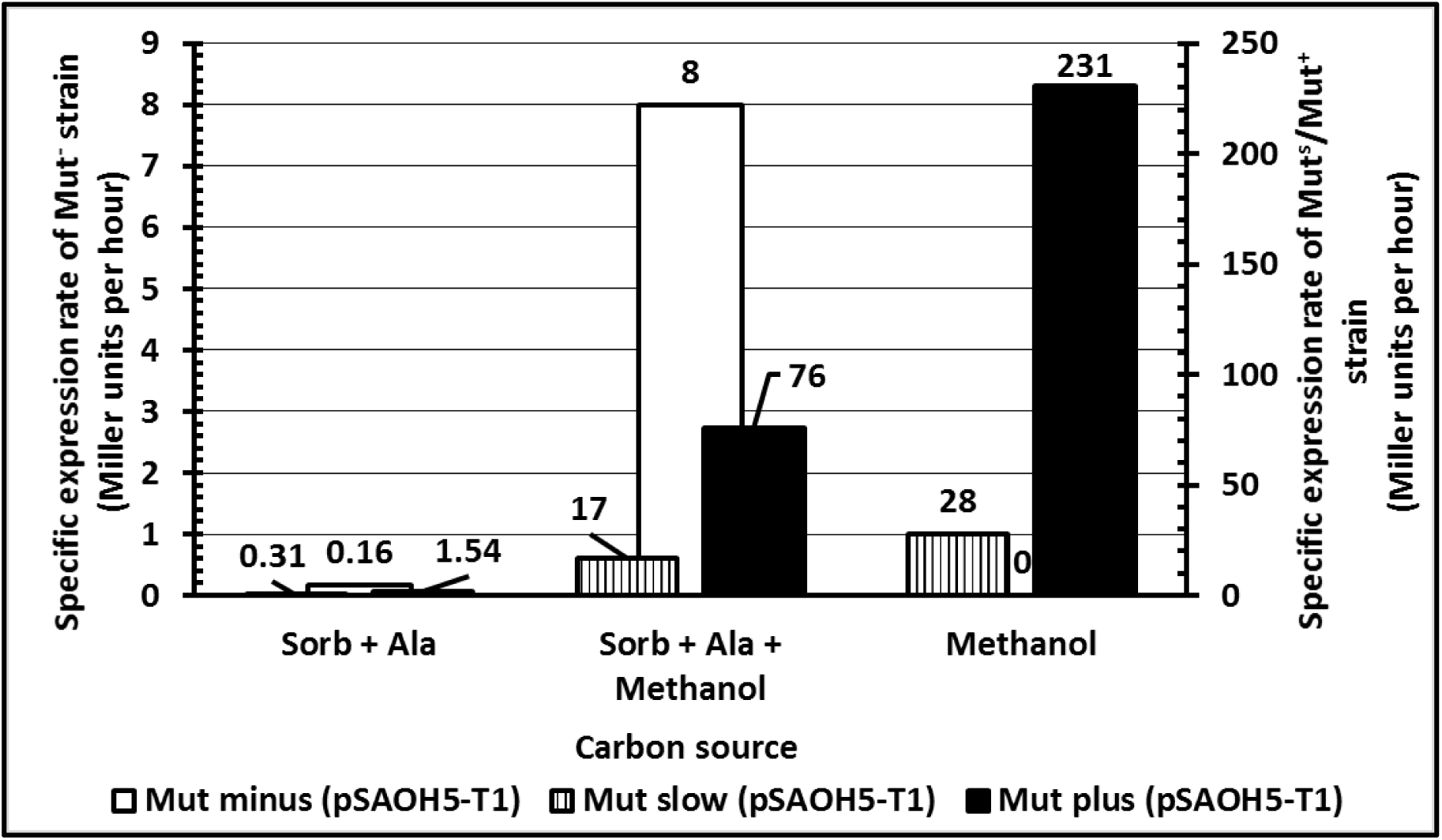
The specific β-galactosidase activity and specific expression rate in Mut^+^ strains is 5-10 times than that observed in Mut^s^ and Mut^−^ strains. Strains Mut^+^ (pSAOH5-T1), Mut^s^ (pSAOH5-T1) and Mut^−^ (pSAOH5-T1) were grown on minimal medium supplemented with either a mixture of sorbitol (55 mM) and alanine (112 mM) with (at various concentrations between 0-160 mM)/ without methanol or methanol alone (160 mM) as a carbon source. Culture samples were collected in the exponential phase when the OD_600_ was between 1-2, and the specific β-galactosidase activities were measured using the modified Miller assay (Materials and Methods). The specific activities and specific growth rates were then plotted against methanol concentration to obtain the maximal expression rates of each Mut strain (Fig. S4a). Specific expression rate at a particular concentration was calculated by multiplying specific β-galactosidase activity with specific growth rate for that particular concentration, as shown in Eq. 3.

To preclude the possibility that the data obtained under inducing conditions is an artefact stemming from the presence of sorbitol + alanine in the medium, we also compared the steady state expression rates of the Mut^+^ and Mut^s^ strains in the presence of pure methanol (160 mM). Under this condition, the specific activities of the Mut^+^ and Mut^s^ strains were similar (2100 and 2800 Miller units, respectively), but the specific growth rate of the Mut^s^ strain (0.01 h^−1^) was substantially lower than that of the Mut^+^ strain (0.11 h^−1^). Consequently, Eq. (3) implies that the specific expression rates of the Mut^+^ and Mut^s^ strains are 231 MU h^−1^ and 28 MU h^−1^, respectively. Thus, the specific expression rates of Mut^+^ and Mut^s^ strains in the presence of pure methanol are 3 and 1.5 times those obtained in the presence of sorbitol + alanine + methanol, which is presumably due to the lower methanol consumption rate in the latter case. However, even in the presence of pure methanol, the specific expression rate of the Mut^+^ strain is 8 times that of the Mut^s^ strain.

It is therefore clear that the specific β-galactosidase expression rates observed under inducing conditions are precisely the opposite of the trends predicted by the model. The model predicts that if methanol is transported by diffusion and the sole inducer of AOX expression, then the specific expression rates under inducing conditions must follow the trend Mut^+^ < Mut^s^ < Mut^−^ (Fig. 1), but the observed rates follow the opposite trend Mut^+^ » Mut^s^ > Mut^−^ (Fig. 2). We conclude that at least one of the two assumptions of the model is untenable and our next goal is to determine if the first assumption of diffusive transport is valid.

### 2.3 Methanol is transported by diffusion

As noted earlier, there are no published reports on the kinetics and mechanism of methanol transport in methylotrophic yeasts, but it is implicitly assumed that methanol is transported by diffusion. We decided to verify this by studying the uptake of ^14^C-methanol.

To determine the kinetics and mechanism of methanol transport, it is convenient to work with a strain that cannot metabolize methanol. If such a strain is exposed to methanol, the intracellular methanol concentration increases until steady state is reached, and transport is diffusive if the steady state concentration of intracellular methanol is essentially equal to the concentration of extracellular methanol. It is necessary to perform this experiment with both uninduced and induced cells, since active transport, even if it exists, may not occur in uninduced cells.

Therefore, methanol uptake kinetics were determined and compared in both uninduced as well as induced cells of the Mut^−^ strain. Briefly, cells were grown in a sorbitol-alanine-based minimal medium with or without methanol, harvested during the exponential phase, and then exposed to ^14^C-methanol (3 mM, 4 x 10^7^ cpm/ml). Culture samples were harvested at regular intervals and measured for radioactive counts. Intracellular methanol concentration was determined by assuming that the volume of a *K. phaffii* cell is ∼3 x 10^−11^ ml (Kunert *et al.*, 2008) and 68 % of cell volume is occupied by water (Kamihira *et al*., 1987).

We found the same uptake kinetics and steady state intracellular ^14^C-methanol levels in both uninduced and induced cells (Fig. 3). The estimated steady state intracellular methanol concentration was essentially identical to the extracellular methanol concentration (Fig. 3). It follows that even in induced cells, there is no evidence of active transport, and methanol is transported by diffusion.

**Fig. 3:**
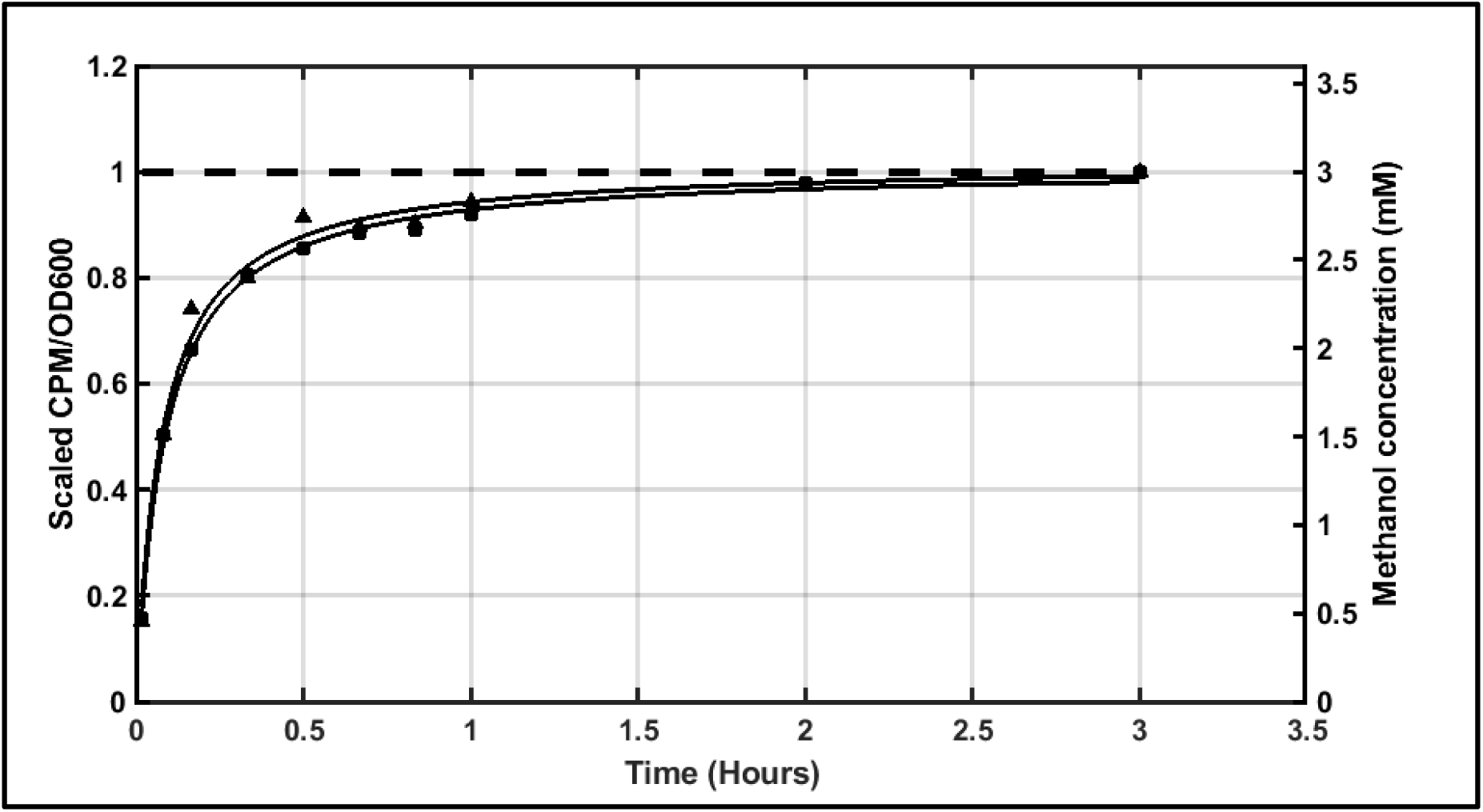
Methanol is transported by diffusion. Strain MC100-3 was grown on either sorbitol (55 mM)/alanine (112 mM) (Uninduced) or sorbitol (55 mM)/alanine (112 mM) plus methanol (160 mM) (Induced) supplemented minimal medium. Cells were harvested in exponential phase (OD_600_ = 0.5-1) and washed. At t = 0, radiolabelled methanol was added to both the cultures. Samples were collected at particular time intervals and immediately filtered through a membrane filter (pore size 0.45 micron). Intracellular counts were measured after washing the filters and dissolving them in a scintillation cocktail (Materials and Methods). The dotted line represents the extracellular concentration of methanol added to the medium. Intracellular concentration was calculated by using 3 x 10^−11^ ml as cell volume (Kunert *et al.*, 2008) and 68 % water content per cell (Kamihira *et al*., 1987).

In light of these observations, we conclude that the second assumption of our model, namely methanol is the sole inducer of P_*AOX1*_, is untenable.

### 2.4 Different Mut^+^, Mut^s^ and Mut^−^ expression rates are due to differences in AOX activity

We have shown above that the specific expression rates follow the trend Mut^+^ » Mut^s^ > Mut^−^, which is the opposite of the trend Mut^+^ < Mut^s^ < Mut^−^ implied by the model. Now the fundamental difference between a Mut^+^ and a Mut^s^/Mut^−^ strain is the presence or absence of AOX1 activity. Mut^+^ strains have a functional copy of the *AOX1* gene, whereas a large fragment of this gene is deleted from Mut^s^/Mut^−^ strains (Cregg *et al*., 1989). It is therefore likely that the different expression rates of Mut^−^, Mut^s^, and Mut^+^ strains are due to differences in their AOX activities. However, to eliminate the possibility that the different expression rates were due to any other cause, we decided to verify if:

a. Expression of AOX in Mut^−^ cells enhances the expression level to that characteristic of Mut^+^ cells.
b. Deletion of AOX in Mut^+^ cells reduces the expression levels to that characteristic of Mut^−^ cells.

#### a) Expression of AOX in Mut^−^ (pSAOH5-T1)

To express *AOX* in the strain Mut^−^ (pSAOH5-T1), a plasmid vector with a copy of the *AOX1* gene (pSC-X1) (Materials and Methods) was transformed into strain Mut^−^ (pSAOH5-T1). The transformants were selected on the basis of growth on methanol and/or zeocin (the plasmid used for transformation has a copy of zeocin resistance gene), as expression of AOX1 enzyme should allow Mut^−^ (pSAOH5-T1) cells to grow on methanol. β-galactosidase assays were performed on the selected transformants after growing them in inducing-and uninducing-medium. It was observed that expression of AOX1 in strain Mut^−^ (pSAOH5-T1) increased the expression of β-galactosidase by 5-fold (see Mut^−^ (T1X1) in Fig. 4).

**Fig. 4:**
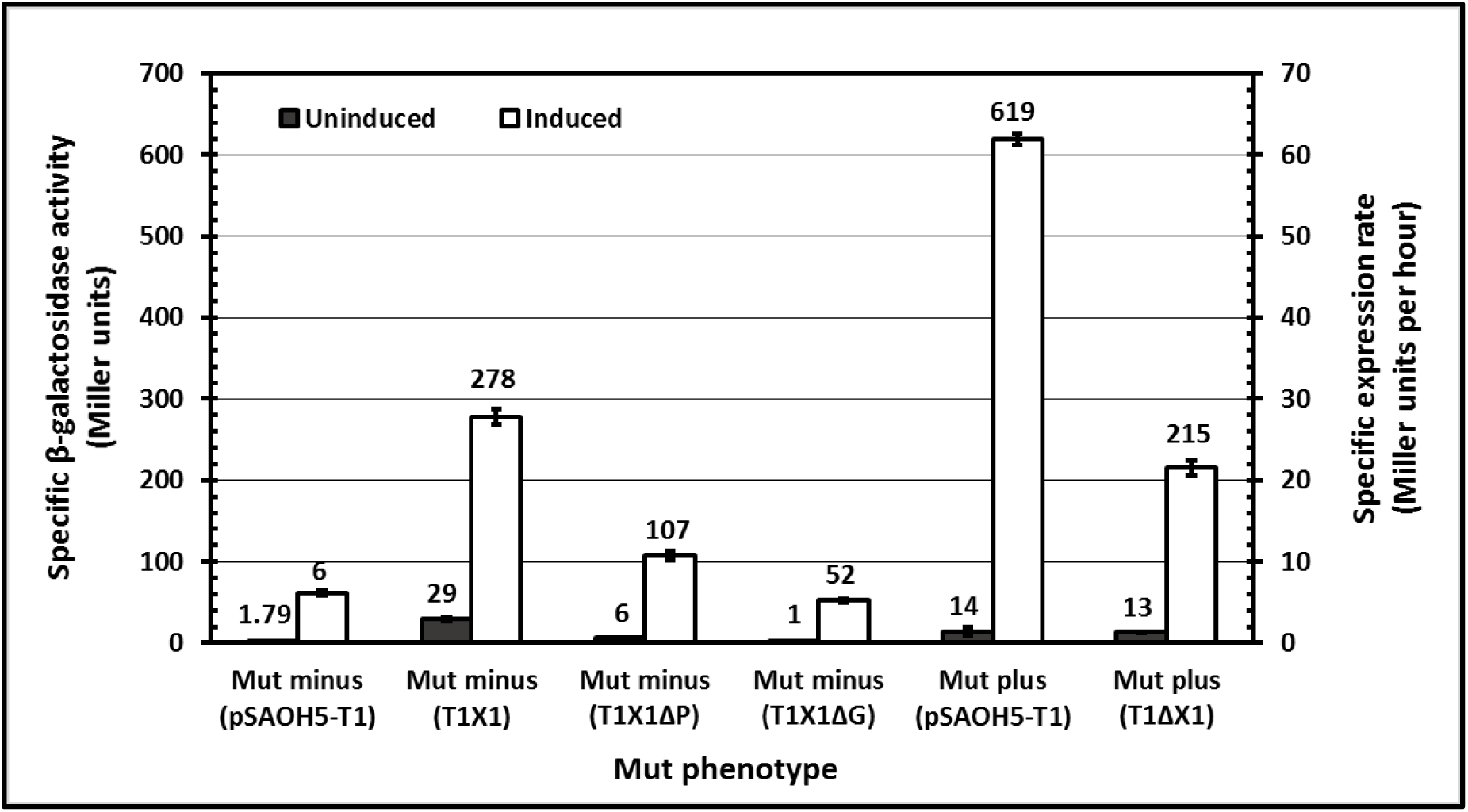
Different expression rates of Mut strains are due to their AOX activities. Cultures were grown on either sorbitol (55 mM) + alanine (112 mM) (Uninduced) or sorbitol (55 mM) + alanine (112 mM) + methanol (160 mM) (Induced) based minimal medium. Samples were collected in the exponential phase (OD_600_ = 1–2) and β-galactosidase activities were measured using Miller assay (Materials and Methods). Standard errors were calculated using biological replicates of a single transformant. Specific expression rate was calculated by multiplying specific β-galactosidase activity with specific growth rate.

We also performed experiments to confirm if the foregoing increase in β-galactosidase expression was reversible by re-abolishing the AOX activity, which proves that the effect is not just due to the presence of the plasmid. AOX expression was knocked-out by engineering a frame-shift mutation in the *AOX1* gene via deletion of an enzyme site (Mut^−^ (T1X1ΔG)) or by deleting the *AOX1* promoter (Mut^−^ (T1X1ΔP)) (for details, refer to Materials and Methods). As predicted, β-galactosidase expression decreased in both cases (Fig. 4).

#### b) Deletion of AOX gene from Mut^+^ (pSAOH5-T1)

For this purpose, a knock-out cassette containing the upstream and downstream flanking chromosomal sequences of *AOX1* gene along with a *Sh ble* gene was constructed (Materials and Methods). In accordance with our expectations, a decline in specific β-galactosidase activity was observed, when the *AOX1* gene was knocked-out from strain Mut^+^ (pSAOH5-T1) (See Mut^+^ (T1ΔX1) in Fig. 4).

AOX activities were measured to confirm the phenotypes of the constructed strains (Table S1). Taken together, these experiments confirm that the different expression rates in strains Mut^+^, Mut^s^, and Mut^−^ are due to the differences in their AOX activities.

### 2.5 Formate and formaldehyde induce P*AOX1* expression

The above data show that methanol is transported by diffusion and the P_*AOX1*_ expression rate increases with AOX activity of the strain. This implies that methanol is not the sole inducer, and metabolites downstream of the AOX-catalysed reaction, namely, formaldehyde or formate, also induce P_*AOX1*_ expression. To test this hypothesis, we grew the Mut^+^, Mut^s^, and Mut^−^ strains on mixtures of sorbitol, alanine, and various concentrations of formate or formaldehyde, measured the steady state specific growth rates (Fig. S2b and S2c) and specific β-galactosidase activities (Fig. S3b and S3c), and calculated the corresponding specific β-galactosidase expression rates from Eq. (3).

In the case of formate, the specific expression rates of all three strains plateaued when the formate concentrations were 15–30 mM (Fig. S4b). Figure 5 shows the specific expression rates for all three strains under fully inducing conditions 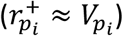. It is clear that formate induces Mut^+^ expression almost as well as methanol. However, this does not imply that formate is, by itself, an inducer of P_*AOX1*_ expression since it is possible that the sole inducer is methanol which is produced in the presence of formate by reversed operation of the methanol dissimilatory pathway. This possibility is ruled out by the data for Mut^s^ and Mut^−^ strains. Indeed, if methanol were the sole inducer of P_*AOX1*_, the Mut^s^ and Mut^−^ strains should display little or no P_*AOX1*_ expression in the presence of formate. Yet, we found that formate induces the Mut^s^ and Mut^−^ strains quite well (Fig. 5). It follows that formate or/and formaldehyde, by themselves, induce P_AOX1_ expression.

**Fig. 5:**
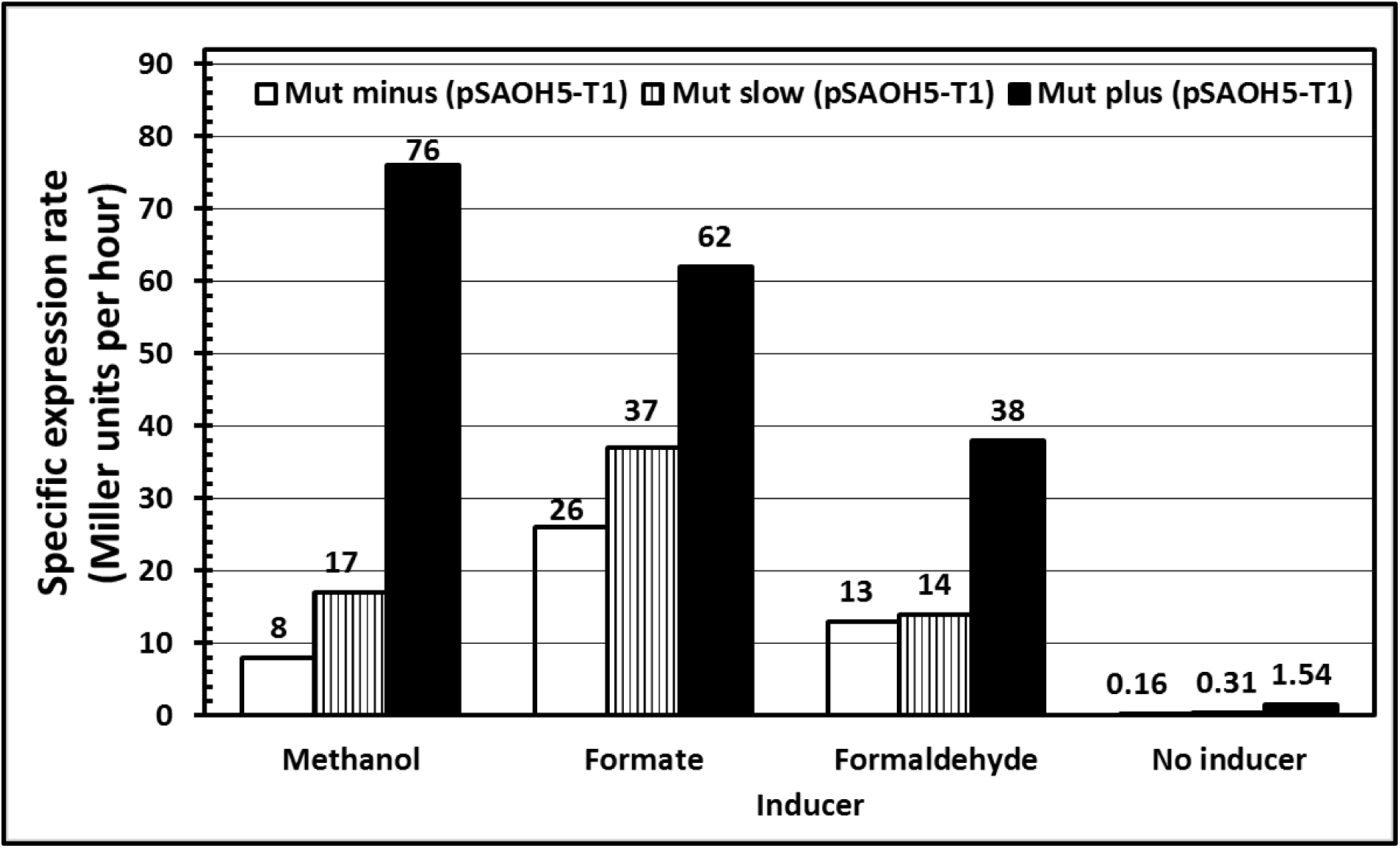
Comparison of methanol, formate and formaldehyde as inducers of protein expression in the three Mut phenotypes. Strains Mut^+^ (pSAOH5-T1), Mut^s^ (pSAOH5-T1) and Mut^−^ (pSAOH5-T1) were grown on a mixture of sorbitol (55 mM) and alanine (112 mM) with either methanol (0-160 mM), potassium formate (0-60 mM) or formaldehyde (0-2 mM). Culture samples were collected in the exponential phase (OD_600_ = 1–2) and β-galactosidase activities were measured using Miller assays (Materials and Methods). Specific expression rate at a particular concentration was calculated by multiplying specific β-galactosidase activity with specific growth rate at that concentration (Eq. 3). Maximum specific expression rates achievable (In case of methanol and formate, it was the obtained by fitting the data to a saturating hyperbolic function (Fig. S4a and S4b). However, in case of formaldehyde, the data could not be collected in the saturation regime as higher concentration of formaldehyde were toxic for growth. Therefore, expression rates at 2 mM have been plotted in the figure (Fig. S4c)) with the three inducers or no inducer have been shown in the figure.

In the case of formaldehyde, we were unable to determine the specific expression rates under fully inducing conditions because even in the presence of relatively low formaldehyde concentrations (3 mM), the growth curves of all three strains exhibited extremely long lags of ∼1 week, which was probably due to substrate inhibition (Andrews, 1968). However, we are led to conclude once again that formaldehyde or/and formate are potent inducers of P_AOX1_ expression since the expression rate of all three strains increased dramatically even at the low formaldehyde concentrations used in our work (Fig. S4c).

Since methanol induces the Mut^−^ strain rather poorly when compared to formate (Fig. 5), there is the distinct possibility that methanol is transported actively, but our transport assay did not reveal this because we used methanol as the inducer which failed to induce synthesis of the active transporter sufficiently (Fig. 3). To exclude this possibility, we studied methanol uptake in Mut^−^ cells induced with formate. For this purpose, cells growing exponentially on a mixture of sorbitol and alanine without or with formate (30 mM) were harvested, washed, suspended in a medium containing radio-labelled methanol but no formate, and processed as described earlier. Again, uninduced and induced cells yielded the same methanol uptake kinetics and steady state intracellular methanol concentrations, and the latter were essentially equal to the extracellular methanol concentration (Fig. S5). This confirms that uptake of methanol does not involve any inducible component and occurs via diffusion.

## 3. Discussion

We have shown above that in the presence of methanol, the P_*AOX1*_ expression rate of Mut^+^ is 5-and 10- fold higher than that of the Mut^s^ and Mut^−^ strains, and this occurs because formaldehyde or/and formate, by themselves are potent inducers of P_*AOX1*_ expression. Both results contradict the prevailing consensus which maintains that expression in Mut^+^ is inferior to that in Mut^−^ and Mut^s^, and methanol is the sole inducer. In the first two sections below, we re-examine the literature to identify potential sources of the discrepancies. In the last section, we shall discuss if the use of formate and formaldehyde as inducers can overcome the known limitations of methanol.

### Mut^+^ expresses recombinant proteins much faster than Mut^s^ and Mut^−^

Before re-examining the data in the literature, we reiterate that the key variable of our interest is the specific expression rate of P_*AOX1*_, which, as shown by Eq. (1), depends on several rates.

In our experiments, β-galactosidase was not accumulated, degraded, and secreted which implied that its specific expression rate could be estimated from Eq. (3). It turns out that the degradation rate was not measured in any of the studies listed in Table 2. We have therefore analysed their data by neglecting degradation 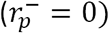 0.

We begin by analysing the studies in which the recombinant proteins were not secreted 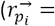. Cregg *et al.* compared the expression of Hepatitis B surface antigen (HBsAg) in steady state Mut^+^ and Mut^s^ cultures (*dp*_*i*_/*dt* = 0) growing exponentially in shake flasks containing 0.5 % w/v methanol (Cregg *et al.*, 1987). They noted in the abstract that “the highest level of HBsAg is observed in a novel mutant [Mut^s^] strain,” but closer inspection reveals that the authors are referring to HBsAg 22 nm particles assembled from HBsAg and lipids. Insofar as the HBsAg protein *per se* is concerned, “quantitative immunoblots indicated that the concentration of HBsAg protein in GS115 (pTBO-6) [Mut^+^ strain] was similar to that in GS115 (pBSAISI) [Mut^s^ strain].” Since the Mut^+^ strain grows 5–30 times faster than the Mut^s^ strains (Tschopp *et al*., 1987b, Brierley *et al*., 1990), Eq. (3) implies that the specific productivity of the Mut^+^ strain is 5–30 times that of the Mut^s^ strain. This may account for the poor assembly of HBsAg 22 nm particles in the Mut^+^ strain since the particles are likely to assemble properly only if synthesis of HBsAg and lipid are coordinated, but this does not occur when HBsAg is synthesized rapidly in the Mut^+^ strain. Clare *et al.* obtained similar results when they compared the expression of tetanus toxin fragment C in Mut^+^ and Mut^s^ cultures growing exponentially in shake flasks containing 1 % v/v methanol (Clare *et al.*, 1991). They observed that “in strains containing nine copies of the expression cassette, fragment C levels were about 20 % lower in the Mut^+^ background compared to Mut^s^.” However, in “single copy transformants the expression level in a Mut^+^ background was similar to or slightly higher than Mut^s^.” Eq. (3) implies that the specific productivity of the single copy Mut^+^ strain must be significantly higher than that of the single copy Mut^s^ strain. Our data with pure methanol (Fig. 2) are consistent with these results — the specific β-galactosidase activities of both strains are essentially the same, but the specific growth rate, and hence the specific expression rate, of the Mut^+^ strain is substantially higher.

Next, we consider the studies in which the recombinant protein was secreted. In virtually all these studies, the authors measured the protein secretion kinetics 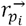, but did not quantify the rates of intracellular protein accumulation *dp*_*i*_/*dt* and dilution *μp*_*i*_(Peng *et al*., 2004, Cos *et al*., 2005, Pla *et al*., 2006, Kim *et al*., 2009, Krainer *et al*., 2012). Under these conditions, Eq. (1) implies that the specific expression rate 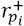 cannot be determined without additional information. In the two studies wherein such information is available, the specific expression rate of the Mut^+^ strain is significantly higher than that of the Mut^s^ strain. Tschopp *et al.* compared the expression of invertase in Mut^+^ and Mut^s^ cultures growing exponentially in shake flasks containing 0.5 % w/v methanol (Tschopp *et al.*, 1987b). They did not report the magnitudes of the terms in Eq. (1), but their pulse-chase experiments showed that the transit time for transport of invertase from the cytosol to the periplasm was 1.5 h in the Mut^+^ strain and 20 h in the Mut^s^ strain. The authors argued that this striking difference was due to the slower secretion rate of Mut^s^ strain, but there is no known reason for differences in the secretion rates of Mut^+^ and Mut^s^ strains. It is conceivable that the dramatically different transit times are due to the slower expression rate of the Mut^s^ strain, a phenomenon for which we have provided evidence in this work. Digan *et al.* and Brierley *et al.* found that when Mut^+^ and Mut^s^ strains were grown in exponential fed-batch cultures with relatively constant methanol concentrations, the accumulation rate of bovine lysozyme *dp*_*e*_/*dt* in the Mut^+^ culture was 6 times that in the Mut^s^ culture. Their immunoblots showed that “essentially no lysozyme remained associated with the insoluble cellular fraction,” i.e., *dp*_*i*_/*dt* ≈ 0 and *μp*_*i*_≈ 0, which implied that the “majority of the lysozyme is secreted,” i.e.,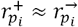. Since both cultures had almost the same biomass concentrations, Eq. (2) implies that the specific expression rate of the Mut^+^ strain must be 6 times faster than that of the Mut^s^ strain assuming negligible protein degradation and dilution (due to volume expansion of the fed-batch culture). In this particular study, the protein secretion rate of the Mut^+^ strain was significantly higher than that of the Mut^s^ strain. However, even if the Mut^+^ and Mut^s^ strains yield comparable protein secretion rates, the expression rate of the Mut^+^ strain may be higher because of the higher protein dilution rate *μp*_*i*_— its faster growth rate allows substantial amounts of protein to be transferred to the new-born cells.

### Formate and formaldehyde induce synthesis of alcohol oxidase

Early studies of several methylotrophic yeasts reported that formaldehyde and formate induced the synthesis of various enzymes of the methanol dissimilatory pathway including alcohol oxidase. Shimizu *et al.* reported that in *Kloeckera* sp., formate (12 g l^−1^) induced synthesis of alcohol oxidase, catalase, formaldehyde dehydrogenase and formate dehydrogenase to levels comparable to those observed with methanol (Shimizu *et al*., 1977). Eggeling *et al.* observed that formaldehyde (3 mM) and sodium formate (70 mM) did not support growth of *Candida boidinii*, but induced expression of the methanol dissimilatory enzymes (Eggeling *et al.*, 1977). Sahm reported the specific activities of the methanol dissimilating enzymes when *C. boidinii* was exposed to various concentrations of methanol, formate and formaldehyde (Sahm, 1977). It was found that these enzymes had the same maximum specific activities regardless of the inducer, but the maximum values were attained at much lower concentrations of formaldehyde (3 mM) than sodium formate (20 mM) and methanol (100 mM).

The above data provided clear evidence that formaldehyde and formate are potent inducers of the methanol dissimilating enzymes, but a subsequent study of an AOX-negative (Mut^−^) strain of *Hansenula polymorpha* led Eggeling and Sahm to conclude that “methanol itself, without being metabolized, acts as inducer on the dissimilatory enzymes” (Eggeling and Sahm, 1980). Specifically, they observed that during growth of an AOX-negative strain on a mixture of glucose and methanol, the specific activities of the methanol dissimilatory enzymes increased 2–3 fold relative to those observed during growth on glucose. These results are consistent with our data — we also found that the expression level of the Mut^−^ strain increased >30-fold upon exposure to methanol (Fig. 2). Assuming that the AOX-negative or Mut^−^ strains do not metabolize *any* methanol, the increase in the dissimilatory enzyme levels must be attributed to induction by methanol itself, i.e., methanol is an inducer of the dissimilatory pathway. However, Eggeling and Sahm concluded that “methanol itself was the effector and not the metabolites derived from it”, i.e., methanol is *the* (sole) inducer of the dissimilatory pathway.

Our data as well as data in the literature, which are summarized below, imply that the methanol cannot be sole inducer of the dissimilatory pathway. Indeed, if methanol is the sole inducer, the P_*AOX1*_ expression rates must be in the order Mut^+^ < Mut^s^ < Mut^−^ (Fig. 1), but we observed exactly the opposite trend Mut^+^ » Mut^s^ > Mut^−^ (Fig. 2). We also found that formaldehyde or/and formate, by themselves, induce P_*AOX1*_ expression since they stimulate expression in the Mut^−^ strain which contains no known pathway from formaldehyde and formate to methanol. Similar results have also been reported in the literature. Tyurin and Kozlov reported that formate induces expression of P_*AOX1*_ in the Mut^+^ strain GS115 of *K. phaffii* and strain Y727*his4* of *K. kurtzmanii* (Tyurin and Kozlov, 2013). Subsequently, they also showed that induction by formate persists in mutants lacking the gene *FLD* encoding formaldehyde dehydrogenase, which implies that induction by formate is not due to FLD-mediated reduction of formate to methanol — instead, formate itself must also an inducer (Tyurin and Kozlov, 2015). Interestingly, they observed that in these FLD mutants, not only formate, but also methanol, induced expression of P_*AOX1*_, which suggests that formate is also an inducer, a result that is consistent with our experiments. They concluded that their experiments “casts doubt on the hypothesis that methanol is the real inducer of the *AOX1* promoter.”

The data presented thus far imply that in addition to methanol, formaldehyde and formate are potent inducers of P_AOX1_ expression, but an alternative hypothesis is conceivable. Indeed, the notion that methanol is an inducer rests on the assumption that the Mut^−^ strain cannot convert *any* methanol to formaldehyde or formate. It is conceivable that formaldehyde or formate are the true inducers, but in the presence of methanol, significant amounts of formaldehyde and formate are produced because a small residual AOX activity persists in the presumably AOX-negative Mut^−^ strain, or there is an AOX-independent pathway from methanol to formaldehyde or formate. This hypothesis explains not only the induction of the Mut^−^ strain in the presence of methanol, but also the dramatically higher expression rates of the Mut^+^ strain relative to the Mut^s^ and Mut^−^ strains which, as noted above, is impossible if methanol is the sole inducer. Further experiments are required to distinguish between the two hypotheses, namely all three metabolites (methanol, formaldehyde, formate) induce or only formaldehyde or/and formate induce.

### The use of formate and formaldehyde to induce recombinant protein expression

Although the chemical identity of the true inducer(s) is unknown, there is clear evidence that formate and formaldehyde are potent inducers of P_*AOX1*_, and therefore promising candidates for recombinant protein production. Yet, we are aware of only two studies in which formate and formaldehyde were used to induce protein production. In the first study, Giuseppin *et al.* reported that both formate and formaldehyde stimulated strong expression of *AOX* in a catalase-free mutant of *H. polymorpha* (Giuseppin *et al*., 1988). Specifically, they observed that when this mutant was grown in a chemostat operated at a dilution rate of 0.1 h^−1^ and fed with various mixtures of glucose and formate or formaldehyde, high specific activities of alcohol oxidase, comparable to those obtained with mixtures of glucose and methanol, were obtained when the molar ratios of formate/glucose and formaldehyde/glucose in the feed were around 2. Later on, Giuseppin *et al.* performed similar studies with a strain of *H. polymorpha* expressing α-galactosidase under the control of the *AOX* promoter, and found that the volumetric productivity obtained (5.5 mg l^−1^ h^−1^) was among the highest reported in the literature (Giuseppin *et al*., 1993). These data confirm that formate and formaldehyde are excellent inducers, and led the authors to note that (Giuseppin *et al*., 1988), “Although methanol is regarded as the actual inducer of the C-1-metabolic pathway (Eggeling *et al.* 1977), more studies are needed to explain the high inducing capacity of formaldehyde and formate.”

Although formate induces as well as methanol, it does not appear to suffer from the limitations that afflict methanol, and it may be useful to study the use of formate (rather than methanol) as inducer of recombinant protein production. Indeed, unlike methanol, formate is non-flammable and hence, relatively safe to use on an industrial scale. Since formate has a much lower heat of combustion (−254 KJ mol^−1^) than methanol (−727 KJ mol^−1^), it will also mitigate other major issues associated with the use of methanol, such as high heat generation and oxygen demand. Finally, formate is an ideal inducer since it supports induction, but very little growth. Hence, addition of formate during the “induction phase” of fed-batch growth will lead to significant recombinant protein, but not biomass, synthesis, thus leading to relatively pure intracellular proteins. Finally, the true potential of Mut^s^ and Mut^−^ strains can only be realized by inducing protein expression with formate (instead of methanol) since these strains would then yield the desired recombinant protein at high levels without simultaneously producing large quantities of the undesired alcohol oxidase.

We found that in our batch cultures, formaldehyde was a potent inducer, but inhibited growth strongly. However, its successful use by Giuseppin *et al.* suggests that despite the toxicity of formaldehyde, it can be used as an inducer provided its feed rate is tightly controlled (Giuseppin *et al*., 1988), as is the case in continuous and fed-batch cultures. Further studies are necessary to investigate the feasibility of formaldehyde as inducer.

## 4. Conclusion

Although the *AOX1* promoter of *K. phaffii* is widely used for heterologous protein production, the mechanisms regulating its expression are not well understood. For instance, it is assumed, without definitive evidence, that methanol is transported by diffusion, and is the sole inducer of P_*AOX1*_ expression. In this work, we demonstrated for the first time that methanol is taken up by diffusion, but methanol is not the sole inducer of P_*AOX1*_ expression. For if the latter were true, the diffusive transport of methanol implies that the Mut^+^, Mut^s^, and Mut^−^ strains would have intracellular methanol concentrations, and hence PAOX1 expression rates, in the order Mut^+^ < Mut^s^ < Mut^−^, but our experiments showed exactly the opposite trend Mut^+^ » Mut^s^ > Mut^−^. We verified that the observed trend is due to differences of the AOX activities because expression (resp., deletion) of *AOX* in the Mut^−^ (resp., Mut^+^) strain transforms the P_*AOX1*_ expression rates to those characteristic of the Mut^+^ (resp., Mut^−^) strain. These results suggest that the products of AOX metabolism, namely formaldehyde or/and formate, also induce P_AOX1_ expression. We confirmed that this is indeed the case because both formaldehyde and formate induced P_AOX1_ expression in the Mut^−^ strain which possesses no known pathway from formaldehyde and formate to methanol. We argue that formaldehyde and formate are useful alternatives to methanol since they do not suffer from the deficiencies that afflict methanol.

## 5. Materials and Methods

### 5.1 Strains and plasmids

*K. phaffii* strains GS115 (*his4*), KM71 (*aox1Δ::SARG4 his4 arg4*) and MC100-3 (*aox1Δ::SARG4 aox2Δ::Phis4 his4 arg4*) (also referred to as strains Mut^+^, Mut^s^ and Mut^−^, respectively) (Table 1) and plasmid pSAOH-5 (Tschopp *et al.*, 1987a) (Fig. S6) were provided by J. M. Cregg, Keck Graduate Institute, Claremont, CA, USA. Phenotypes were confirmed by growing the strains on appropriate selective plates. Strain DH5α (Hanahan, 1983) of *Escherichia coli* was used for all cloning and plasmid propagation steps.

### 5.2 Inoculum preparation and culture conditions

Lysogeny broth (LB) and Lysogeny agar (LA) plates were used for growing *E. coli* cells, while *K. phaffii* cells were either grown in a rich YPD medium (1 % yeast extract, 2 % peptone and 1 % glucose) or a buffered minimal medium containing 100 mmoles potassium phosphate buffer (pH 5.5), 15.26 g NH_4_Cl, 1.18 g MgSO_4_.7H_2_O, 110 mg CaCl_2_.2H_2_O, 2.4 mg biotin and 10 ml trace element stock solution (45.61 mg FeCl_3_, 28 mg MnSO_4_.H_2_O, 44 mg ZnSO_4_.7H_2_O, 8 mg CuSO_4_.5H_2_O, 8.57 mg CoCl_2_.6H_2_O, 6 mg Na_2_MoO_4_.2H_2_O, 8 mg H_3_BO_3_, 1.2 mg KI, 370 mg disodium EDTA (ethylene diamine tetra acetic acid) (pH 8.0)) per liter (adapted from Jungo *et al.*, 2006) with an appropriate carbon source (concentrations used are mentioned in the results section). All medium components were prepared as stock solutions, sterilized separately (either by autoclaving or by membrane filtration) and mixed in appropriate volumes before use.

Stock cultures were prepared in medium containing 200 – 300 g l^−1^ glycerol and stored at – 80 °C. Cells were revived by inoculating either YPD medium or minimal medium with the prepared glycerol stocks (primary culture). They were further passaged and allowed to go through at least 5–6 doublings, in order to achieve steady state conditions. Shake-flask cultivations of *K. phaffii* and *E. coli* were carried out at 30 °C and 37 °C, respectively. To maintain aerobic conditions, cultures were grown in vessels with sufficient head-space and were shaken at 250 rpm in a rotary incubator-shaker.

Whenever required, zeocin (Invitrogen) was added to a final concentration of 25 μg ml^−1^ to LB medium for selection in *E. coli.* For selection of resistant *K. phaffii* cells, a final concentration of 50 μg ml^−1^ was used for growth in YPD medium, but a much higher concentration (2 mg ml^−1^) was needed in case of minimal medium. This is due to the high ionic strength of the minimal medium, which is known to inhibit the activity of zeocin. We had to use a much higher concentration than the recommended value of 100 μg ml^−1^, because of the presence of trace elements in our minimal medium recipe.

### 5.3 Construction of a non-autonomous integrative plasmid: pSAOH5-T1

pSAOH-5 is a plasmid that contains an in-frame fusion of the *lacZ* gene (starting with amino acid nine) to the first 15 amino acids of the *AOX1* gene under control of the 5’ regulatory *AOX1* sequence (Tschopp *et al*., 1987a) (Fig. S6). The plasmid also contains the *K. phaffii HIS4* gene under control of its own promoter and thus can complement a *his4* auxotrophy. In addition, plasmid pSAOH-5 contains an ARS (autonomously replicating sequence) for autonomous replication. Some authors have reported that the presence of an ARS sequence makes plasmids unsuitable for stable chromosomal integration, as such a plasmid can integrate at one or more homologous sites (Sreekrishna *et al*., 1997). A self-replicating plasmid can also excise from its integration site (due to the duplication of flanking sequences in case of single crossover recombination) and then can reside in the cell autonomously. This can increase the copy number of the plasmid, which would be undesirable for the purpose of our study. We verified these observations in strains which were transformed with linearized pSAOH-5 by checking for the presence of circular plasmid. Circular full-length plasmid DNA was actually retrieved, when DH5α cells were transformed with a total DNA preparation of a pSAOH-5 containing strain.

To avoid this problem, plasmid pSAOH-5 was modified to remove the ARS fragment and, at the same time, to introduce a transcription terminator region downstream of the *lacZ* gene. Terminator sequences were introduced to stop transcription initiated at the *AOX1* promoter, which is known to be a very strong promoter that might interfere with the expression of a downstream gene, in this case, *HIS4*. The resulting plasmid was named pSAOH5-T1 (Fig. S7).

### 5.4 Construction of single-copy integration Mut strains

*K. phaffii* has an efficient homologous recombination system that can be employed for successfully inserting a gene of interest into the chromosome. One can choose the site of integration by flanking the gene of interest with DNA sequences homologous to the target locus on the genome. To direct integration of the vector into the genome of *K. phaffii* strains, the plasmid is linearized prior to transformation. The site of linearization determines the locus of integration into the genome. In this study, plasmid was linearized by cutting it within the *AOX1* promoter sequence to direct its integration into the *AOX1* locus of the *K. phaffii* genome.

Contrary to the usual recommendation (Chiruvolu *et al*., 1997; Invitrogen manual, 2010), we found that use of SacI for linearizing the plasmid was not ideal, as it cuts the plasmid at two sites. Instead, the vectors pSAOH-5 and pSAOH5-T1 were linearized within the *AOX1* promoter with the single-cutting PmeI prior to transformation. Vector pSAOH-5 has been integrated into the genome of *K. phaffii* before (Tschopp *et al.*, 1987a; Inan and Meagher, 2001), but in these cases, it was inserted into the *his4* locus, which has been reported to yield unstable integration events (Higgins and Cregg, 1998).

In the present work, plasmids were first amplified in *E. coli* strain DH5α and then purified by the alkaline lysis method (Sambrook and Russell, 2001). For single-copy insertions, 50 - 100 ng of the PmeI*-*linearized plasmid DNA were transformed into either Mut^+^, Mut^s^ or Mut^−^ strains via electroporation, after Cregg and Russell (1998) (Fig. S8). Cells were, after transformation, plated on minimal medium plates (containing a mixture of sorbitol, alanine and methanol, along with X-gal, a colorigenic indicator of β-galactosidase activity).

There are multiple ways in which a linearized plasmid can integrate into the chromosomal DNA during single crossover recombination. It is known that homologous recombination often leads to multiple-copy insertions of a plasmid (Clare *et al*., 1991; Schwarzhans *et al*., 2016). These multiple-copy integrations can occur in various orientations like head-to-head, head-to-tail and tail-to-tail, as shown in Fig. S9. Although it is sometimes preferred to have a multicopy insertion of the gene of interest, which might yield more heterologous protein than single-copy events (Cos *et al.*, 2005; Macauley-Patrick *et al.*, 2005), it is necessary to ensure a single-copy integration of the heterologous gene for comparative quantification. After preparation of genomic DNA (See 5.8), transformants were therefore screened for single-copy insertion, using appropriate PCR analysis.

Combinations of three primers (P1, P2 and P3) were used to test for single integration of the plasmid (Table S2 and Table S3). PCR was performed with genomic DNA preparations of several transformants to obtain the desired clones. A particular band pattern with different primer combinations was used as the confirmation test for transformants containing a single copy of the plasmid (Table S2). The band pattern was checked by running the PCR product on 1% agarose-gels (Fig. S10).

### 5.5 Construction of an autonomous plasmid (pSC-X1) for AOX1 expression in Mut^−^ (pSAOH5-T1)

For expression of the *AOX1* gene in strain Mut^−^ (pSAOH5-T1), a self-replicating plasmid was constructed with the following features: a) Selectable markers: for selection in yeast and bacteria, b) ARS: for autonomous replication of the plasmid in *K. phaffii*, c) Origin of replication (ori): for replication of plasmid in *E. coli*, d) *AOX1* gene under the control of its endogenous promoter P_*AOX1*_ (Fig. S11). pBC-KS (+) (Stratagene, LA Jolla, California, USA), was chosen as the basic cloning vector. It already contains an ori for replication and a chloramphenicol resistance gene for selection in *E. coli*.

Zeocin belongs to the family of bleomycin/phleomycin broad spectrum antibiotics. It is used for selection in eukaryotes, such as yeast. The bleomycin resistance is conferred by the product of the *Sh ble* (*Streptoalloteichus hindustanus* bleomycin) gene. It confers resistance to zeocin by inhibiting the DNA cleavage activity of zeocin. pPICZα-B (Invitrogen) belongs to a family of *K. phaffii* vectors that carry the *Sh ble* gene. A DNA fragment containing the *Sh ble* gene was isolated by cutting pPICZα-B with BamHI and EcoRV. A 1 kb NdeI-HindIII fragment containing the ARS sequence was taken out from pSAOH-5. DNA fragments containing the *AOX1* promoter and gene were amplified by PCR using Vent polymerase (NEB) with primer combinations *AOX1*P-F and *AOX1*P-R for the promoter and *AOX1*G-F and *AOX1*G-R for the gene (Table S3).

For transformation into Mut^−^ (pSAOH5-T1) cells, 0.5 – 1 μg of purified plasmid preparation were added to the electro-competent cells before pulsing the cells. Cells were allowed to recover for 5 – 6 hours at 30 °C before plating them on appropriate selective plates. The resulting strain was named Mut^−^ (T1X1).

### 5.6 Abolishing the AOX activity of strain Mut^−^ (T1X1)

AOX expression was abolished in strain Mut^−^ (T1X1) by either deleting the promoter of the *AOX1* gene or by introducing a frame-shift mutation in the *AOX1* gene. For deleting P_*AOX1*_, the PstI-HindIII fragment containing P_*AOX1*_ was removed from pSC-X1. The resulting plasmid was named pSC-X1ΔP (Fig. S12) and transformed into Mut^−^ (pSAOH5-T1), as described earlier.

A frame-shift mutation in a gene alters its protein product and usually truncates it and renders it non-functional. Therefore, to delete AOX1 expression from Mut^−^ (T1X1), a frame-shift mutation was introduced into the *AOX1* gene by deleting a restriction enzyme site (SalI) within the 5’ end of the gene. The new plasmid was named pSC-X1ΔG (Fig. S13).

### 5.7 Construction of a knock-out cassette for deletion of gene *AOX1* from Mut^+^ (pSAOH5-T1)

A BamHI - EcoRV fragment from pPICZα-B containing the *Sh ble* gene was fused with the *AOX1* promoter sequence at the 5’ end and a 1 kb genomic DNA region downstream to the *AOX1* gene at the 3’ end (Fig. S14A). The 5’ and 3’ fragments were amplified with PCR using genomic DNA preparation from strain GS115 and the primers: *AOX1*P-F and Frg.1P-R for the 5’ homologous region and Frg.3-F and Frg.3-R for the 3’ homologous region (Table S3).

These flanking homologous regions facilitated the replacement of the *AOX1* gene with the *Sh ble* gene via a double crossover recombination event (Fig. S14B). The resulting cassette was inserted between the PstI and SalI sites of a cloning vector and amplified in *E. coli.* The cassette was cut out using the same sites and gel-purified before transformation into strain Mut^+^ (pSAOH5-T1).

### 5.8 Genomic DNA preparation

The standard phenol, chloroform and isoamyl alcohol DNA extraction method for *E. coli* was modified to isolate genomic DNA from *K. phaffii* cells. Briefly, cells were harvested in late-log phase and collected by centrifugation. The pellet was washed with sterile distilled water and resuspended in an appropriate volume of lysis buffer (100 mM Tris–HCl (pH 8.0), 20 g l^−1^ Triton X–100, 10 g l^−1^ sodium dodecyl sulfate, 100 mM NaCl, 10 mM Na_2_EDTA) (Schroder *et al*., 2007). For cell lysis, resuspended cells were vortexed vigorously for one minute and then kept at 65 °C in a water bath for 20 minutes, after which they were again vortexed for one minute. Lysis was followed by extraction with phenol-chloroform-isoamyl alcohol and ethanol precipitation of DNA (Sambrook and Russell, 2001).

### 5.9 Cell permeabilization

Cells were grown in minimal medium supplemented with an appropriate carbon source. Culture samples were collected during exponential phase and immediately placed on ice to arrest metabolic activity. Cells were then collected by centrifugation and resuspended in Z buffer (Miller, 1972) (without β-mercaptoethanol). To permeabilize the cells, this was followed by three cycles of snap-freezing in liquid nitrogen and thawing at 30 °C, after adding 25 μl of 1 % Triton X-100 to 0.5 ml aliquots of resuspended cells (adapted from Kippert, 1995).

### 5.10 β-galactosidase assay

Appropriate dilutions of permeabilized cells were used to set up the assays according to Miller (1972). The specific β-galactosidase activity was calculated with the formula

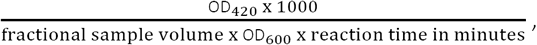

where OD_420_ and OD_600_ represent the optical densities at 420 nm and 600 nm, respectively.

### 5.11 Alcohol oxidase assay

Measurements of alcohol oxidase activity were carried out as described by Jungo *et al.* (2006). Cells were permeabilized using triton and liquid nitrogen as described in section 5.9. A reaction mixture containing 4-aminoantipyrine (AAP), phosphate buffer (pH 7.4), horseradish peroxidase (HRP) and 4-hydroxybenzenesulfonic acid was prepared and added to the permeabilized cells. Reactions were started by addition of methanol. The rate of change in absorbance (500 nm) at 30 °C was determined spectrophotometrically (Spectramax, M2e, Molecular Devices Corp., Sunnyvale, California, USA). Specific AOX activities were calculated using the formula: (Rate of change of OD_500_ per sec x 10^5^)/ (sample volume in ml x OD_600_ of cell suspension used in the assay).

### 5.12 Methanol uptake assay

Cells (Mut^−^) were grown to an OD_600_ of 0.5 - 1.0 in sorbitol plus alanine supplemented minimal medium with (induced) or without methanol/potassium formate (uninduced). Cells were collected by centrifugation at 30 °C and 6000 rpm for 2 min, washed twice with minimal medium (without methanol/potassium formate) and resuspended in the same medium to an OD_600_ of 0.5. ^14^C-methanol (specific activity 57 mCi mmol^−1^) (American Radiolabeled Chemicals, Inc., St. Louis, Missouri, USA) at a final concentration of 2.7 mM was added to the culture (time t = 0) after equilibrating them at 30 °C for 30 minutes. 0.5 ml aliquots were collected at appropriate time intervals and filtered through 0.45 μm nitrocellulose filters. Filters were washed with 3 ml of minimal medium (without methanol) to remove residual extracellular ^14^C-methanol. Filters were then immediately dissolved in 10 ml scintillation cocktail (3 g 2,5-Diphenyloxazole (PPO), 0.2 g 1,4-bis(5-phenyloxazol-2-yl) benzene (POPOP), 257 ml triton X-100, 106 ml ethanol, 37 ml ethylene glycol, 600 ml xylene in 1 liter) (adapted from Fricke, 1975 and Anderson and McClure, 1973) in liquid scintillation vials and counts per minute (cpm) were determined using a liquid scintillation counter (Perkin-Elmer, Boston, Massachusetts, USA).

### 5.13 Software

All the plasmid maps and DNA sequences were created and analysed using Snapgene viewer (GSL biotech; www.snapgene.com). MATLAB R2017b was used for curve fitting application.

