## Supplementary File for "The Mut^+^ strain of *Komagataella phaffii* (*Pichia pastoris*) expresses P*_AOX1_* 5 and 10 times faster than Mut^s^ and Mut^−^ strains: Evidence that formaldehyde or/and formate are true inducers of AOX"

### Supplementary Figures and Tables

#### 1. Supplementary figures

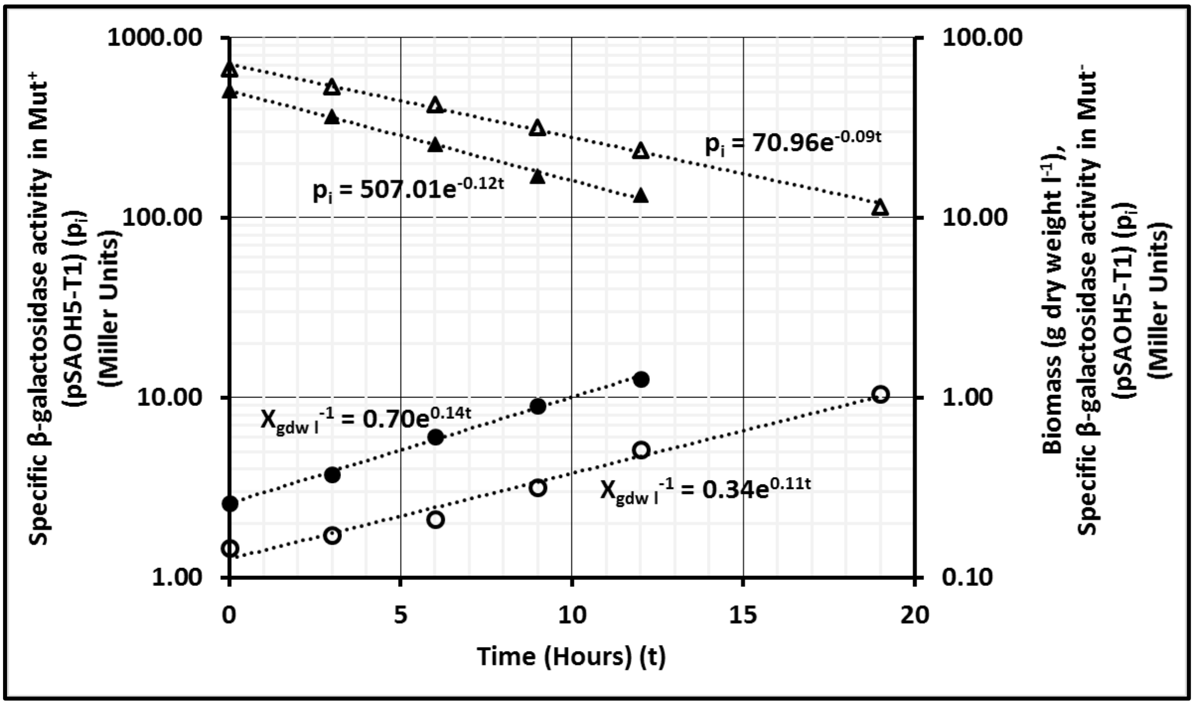


**Fig. S1: The specific β-galactosidase degradation rate in Mut⁺ and Mut⁻ strains.** Mut⁺ and Mut⁻ cultures were grown on minimal medium supplemented with sorbitol (55 mM), alanine (112 mM) and methanol (160 mM) until fully induced. At time (t) = 0 hours, these fully induced cultures were transferred to an inducer free (methanol minus) sorbitol (55 mM) and alanine (112 mM) containing minimal medium. Cell growth was monitored by measuring OD600 of the cultures (Biomass (gdw $l^{-1}$) values were calculated using the expression 1 OD600 = 0.37 g $l^{-1}$). Culture samples were collected to perform β-galactosidase assays as described in Materials and Methods. In the above figure, specific β-galactosidase activities of Mut⁺ (▲) and Mut⁻ (Δ) strains along with biomass values of Mut⁺ (●) and Mut⁻ (○) strains have been plotted against time.

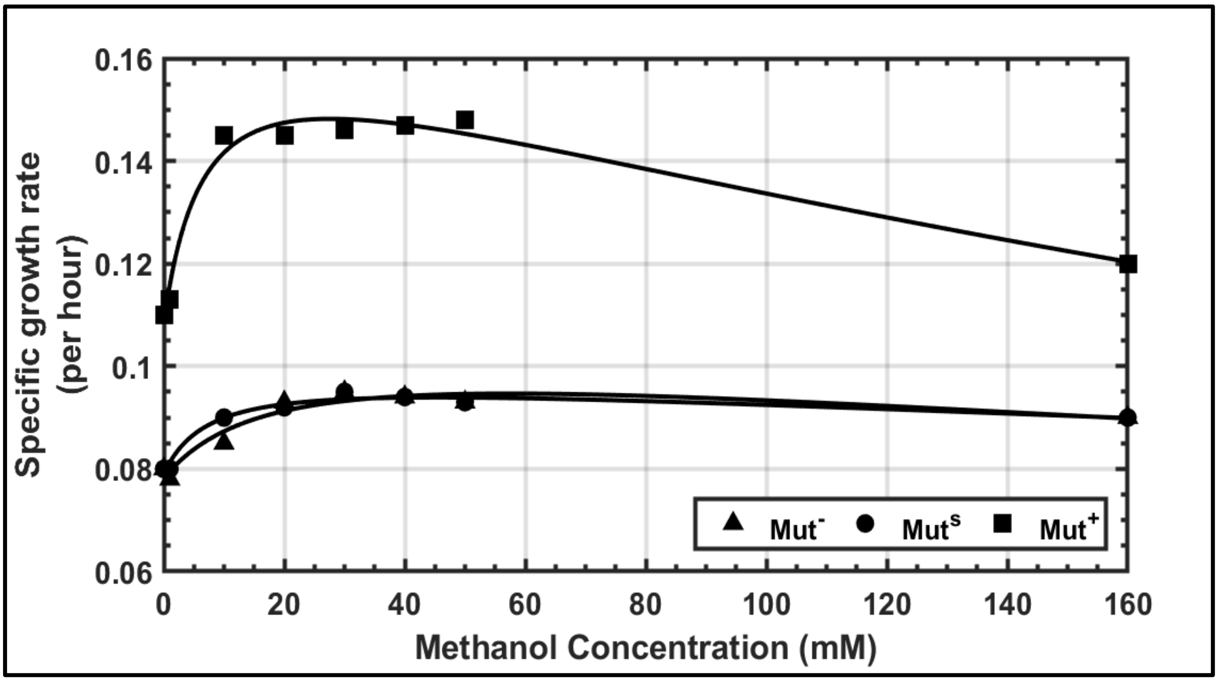


**Fig. S2a: Effect of methanol concentration on specific growth rate of the three Mut strains.** Strains Mut$^+$ (pSAOH5-T1), Mut$^s$ (pSAOH5-T1) and Mut$^-$ (pSAOH5-T1) were grown on minimal medium supplemented with a mixture of sorbitol (55 mM) and alanine (112 mM) with various concentrations (0-160 mM) of methanol as a carbon source. Growth was monitored by measuring OD600 of the cultures at regular time intervals.

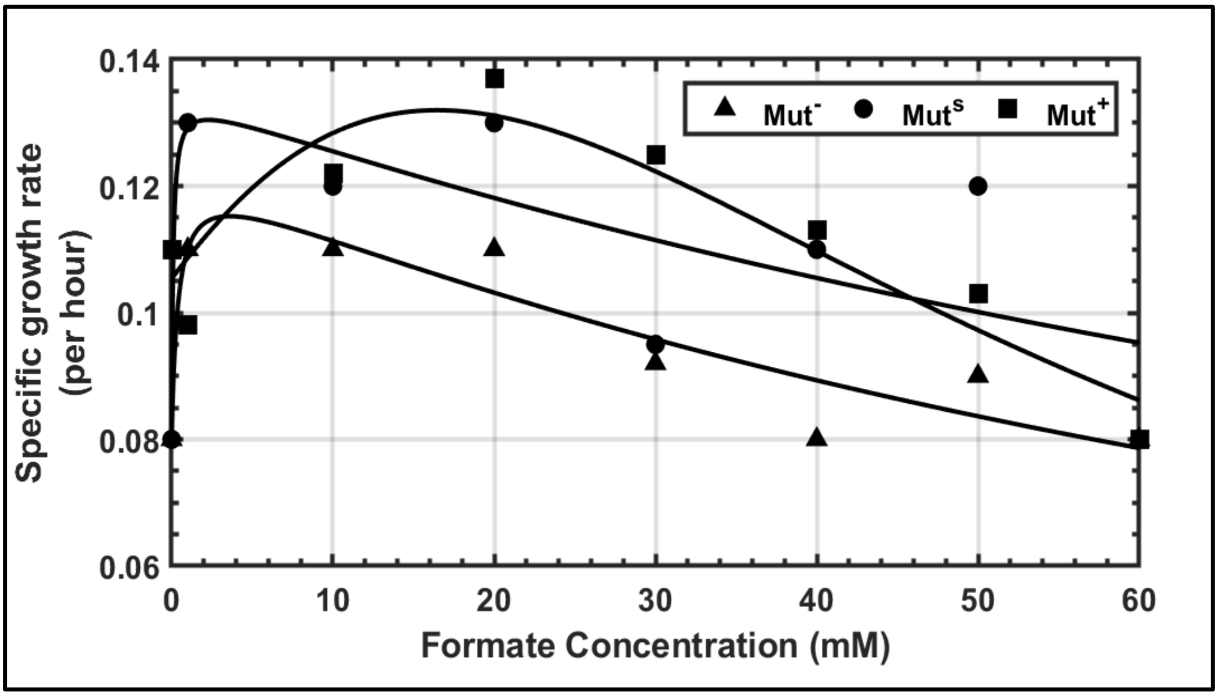


**Fig. S2b: Effect of formate concentration on specific growth rate of the three Mut strains.** Strains Mut$^+$ (pSAOH5-T1), Mut$^s$ (pSAOH5-T1) and Mut$^-$ (pSAOH5-T1) were grown on minimal medium supplemented with a mixture of sorbitol (55 mM) and alanine (112 mM) with various concentrations (0-60 mM) of formate as a carbon source. Growth was monitored by measuring OD600 of the cultures at regular time intervals.

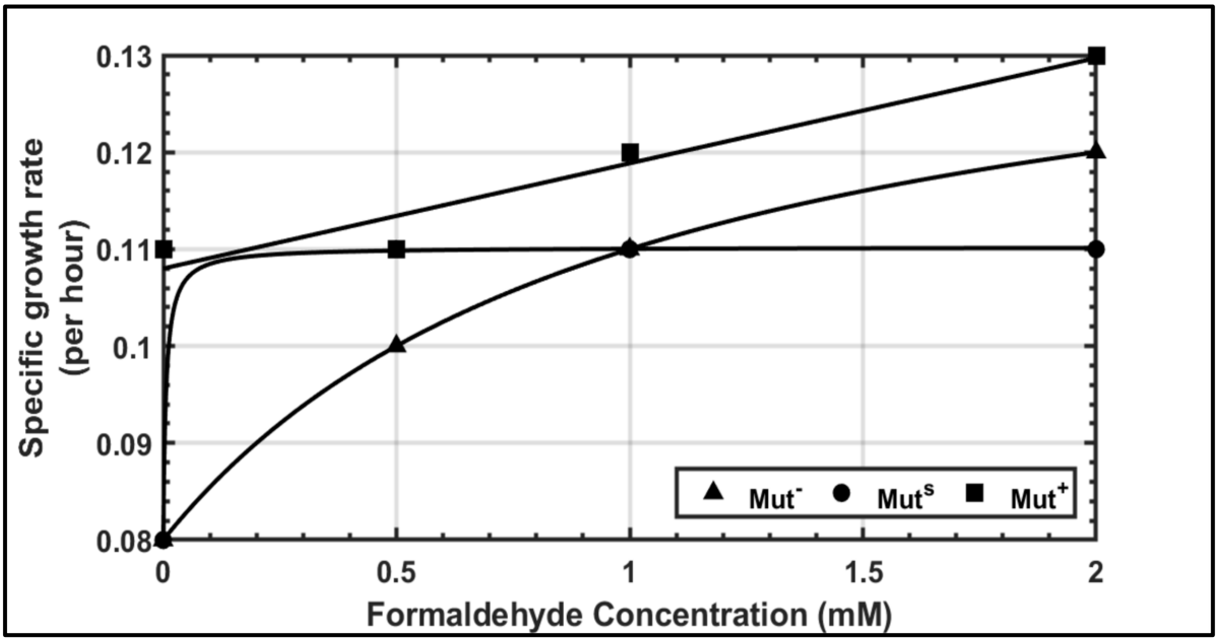


**Fig. S2c: Effect of formaldehyde concentration on specific growth rate of the three Mut strains.** Strains Mut+ (pSAOH5-T1), Mut$^s$ (pSAOH5-T1) and Mut- (pSAOH5-T1) were grown on minimal medium supplemented with a mixture of sorbitol (55 mM) and alanine (112 mM) with various concentrations (0-2 mM) of formaldehyde as a carbon source. Growth was monitored by measuring OD600 of the cultures at regular time intervals.

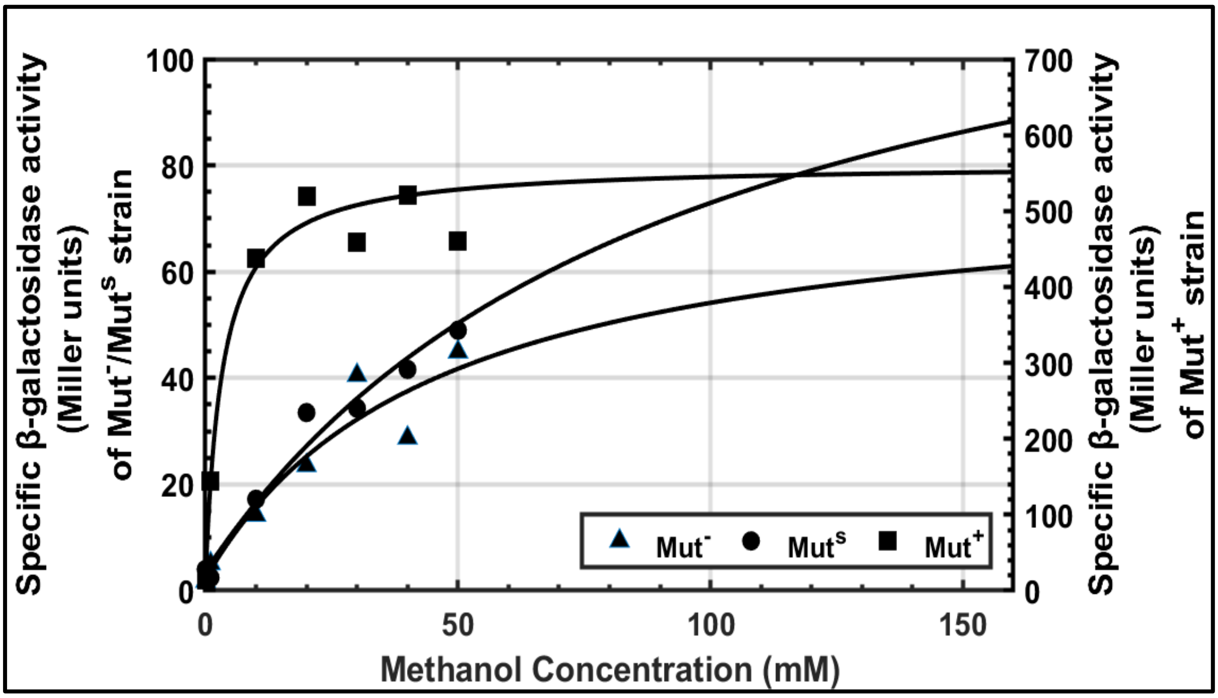


**Fig. S3a: Effect of methanol concentration on specific β-galactosidase activities in the three Mut strains.** Strains Mut+ (pSAOH5-T1), Mut$^s$ (pSAOH5-T1) and Mut- (pSAOH5-T1) were grown on minimal medium supplemented with a mixture of sorbitol (55 mM) and alanine (112 mM) with various concentrations (0-160 mM) of methanol as a carbon source. Culture samples were collected in the exponential phase when the $OD_{600}$ was between 1-2, and the specific β-galactosidase activities were measured using the modified Miller assay (Materials and Methods).

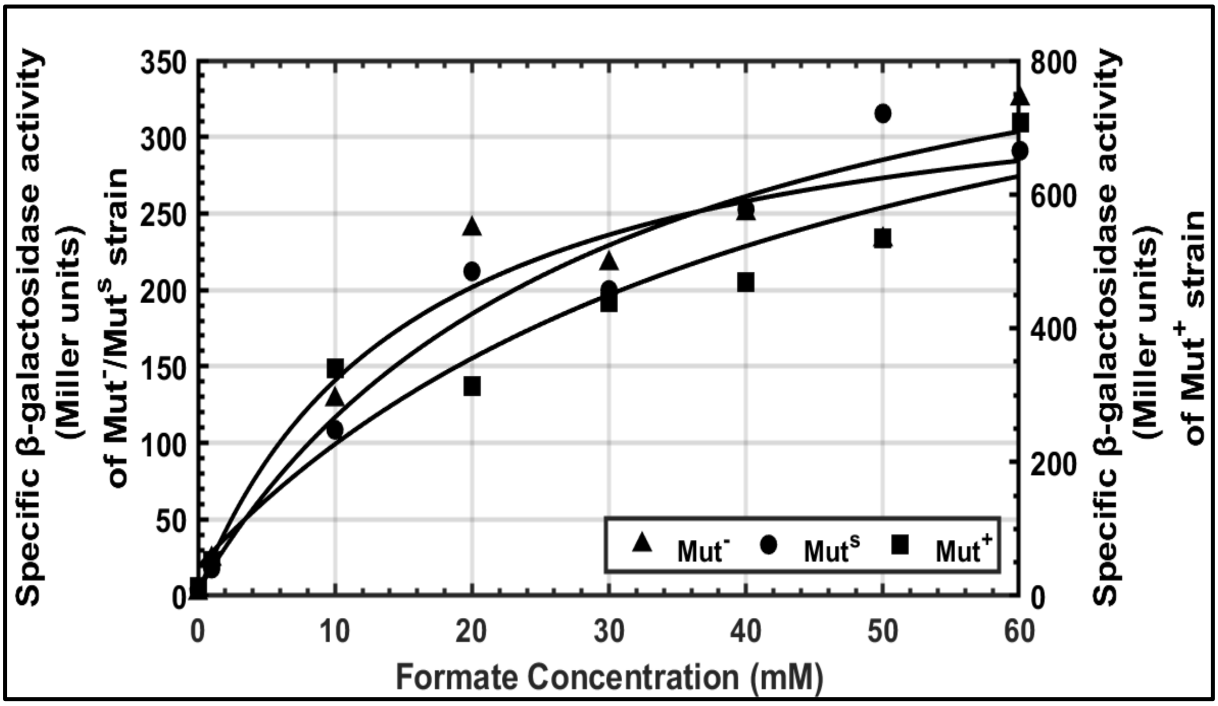


**Fig. S3b: Effect of formate concentration on specific β-galactosidase activities in the three Mut strains.** Strains Mut+ (pSAOH5-T1), Mut^s (pSAOH5-T1) and Mut- (pSAOH5-T1) were grown on minimal medium supplemented with a mixture of sorbitol (55 mM) and alanine (112 mM) with various concentrations (0-60 mM) of potassium formate as a carbon source. Culture samples were collected in the exponential phase when the $OD_{600}$ was between 1-2, and the specific β-galactosidase activities were measured using the modified Miller assay (Materials and Methods).

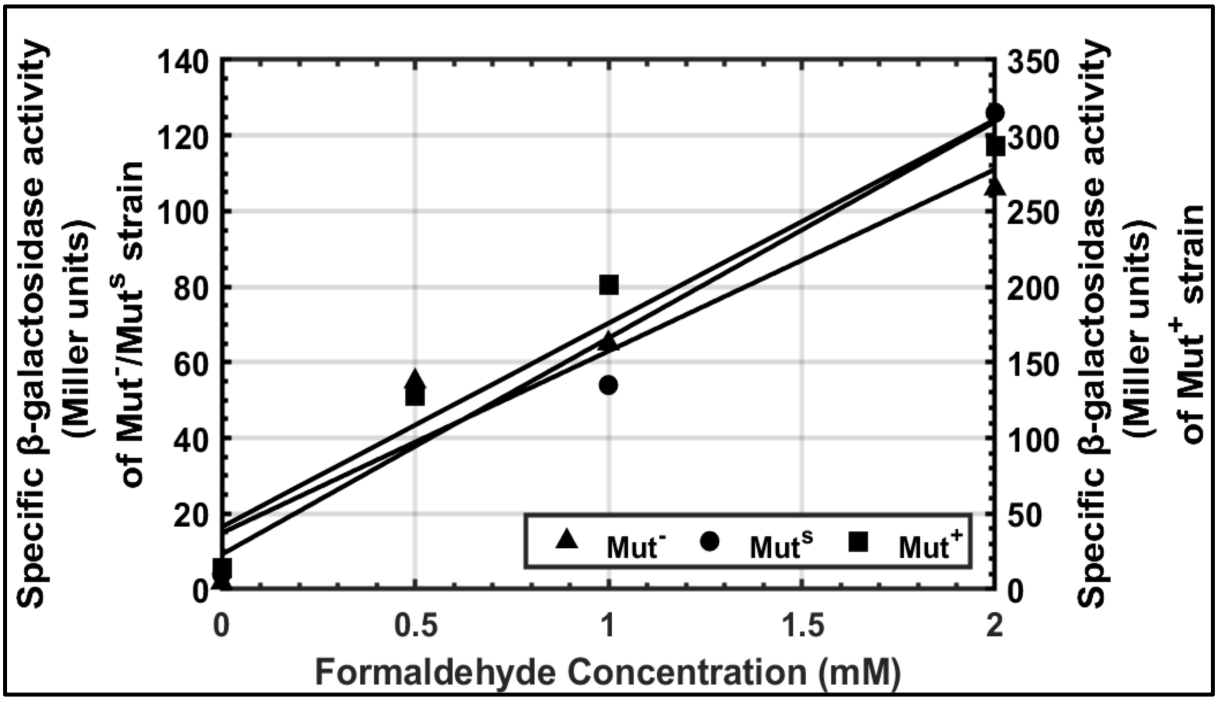


**Fig. S3c: Effect of formaldehyde concentration on specific β-galactosidase activities in the three Mut strains.** Strains Mut+ (pSAOH5-T1), Mut^s (pSAOH5-T1) and Mut- (pSAOH5-T1) were grown on minimal medium supplemented with a mixture of sorbitol (55 mM) and alanine (112 mM) with various concentrations (0-2 mM) of formaldehyde as a carbon source. Culture samples were collected in the exponential phase when the $OD_{600}$ was between 1-2, and the specific β-galactosidase activities were measured using the modified Miller assay (Materials and Methods).

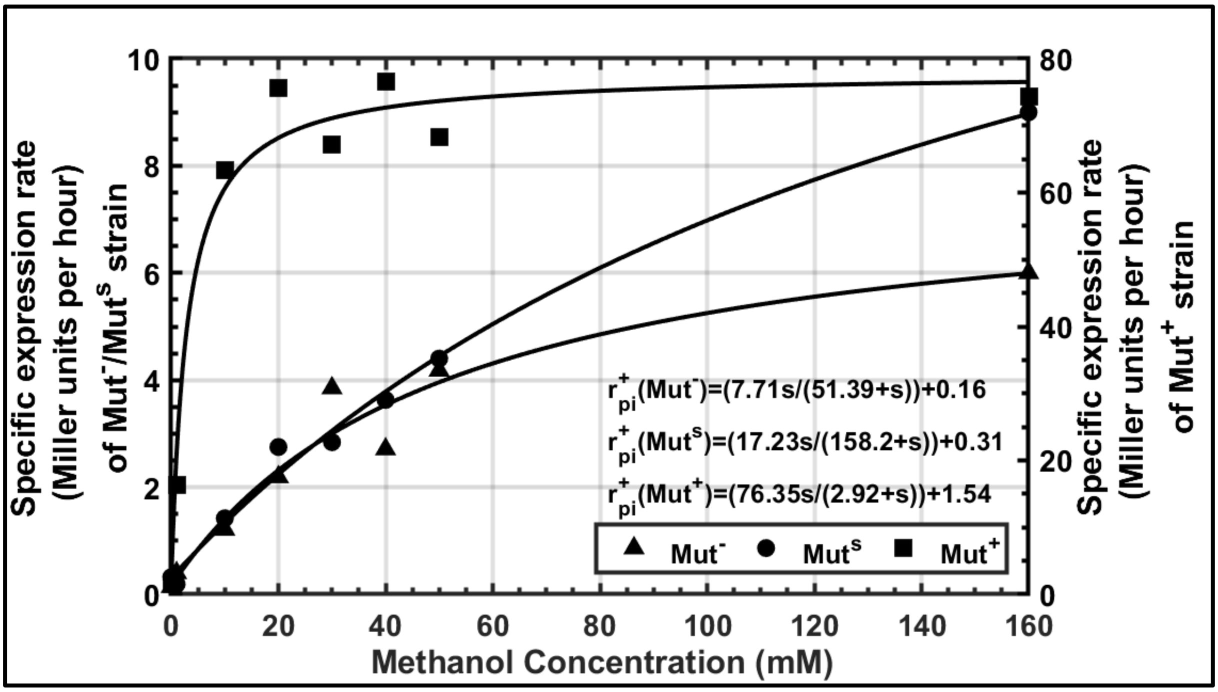


**Fig. S4a: Effect of methanol concentration (s) on the specific expression rate of the three Mut phenotypes.** Strains $Mut^+$ (pSAOH5-T1), $Mut^s$ (pSAOH5-T1) and $Mut^-$ (pSAOH5-T1) were grown on minimal medium supplemented with a mixture of sorbitol (55 mM) and alanine (112 mM) with various concentrations (0-160 mM) of methanol as a carbon source. Culture samples were collected in the exponential phase when the $OD_{600}$ was between 1-2, and the specific β-galactosidase activities were measured using the modified Miller assay (Materials and Methods). Specific expression rate at a particular concentration of methanol was calculated by multiplying specific β-galactosidase activity with specific growth rate at that particular concentration, as shown in Eq. 3.

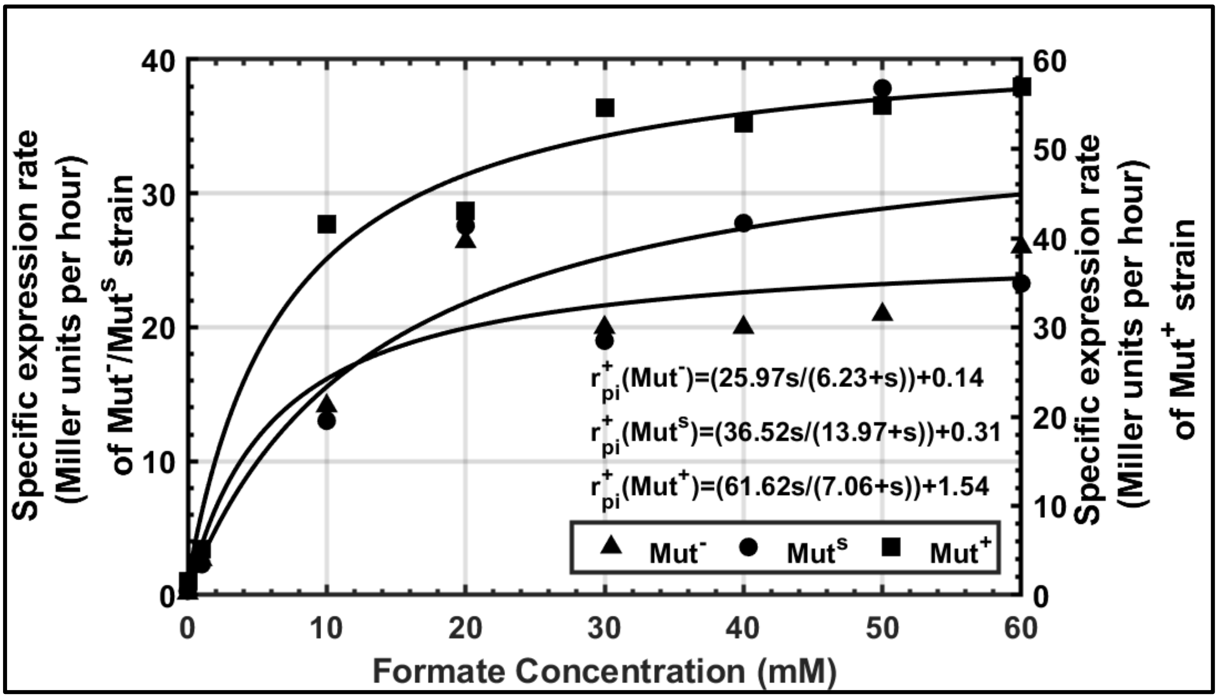


**Fig. S4b: Effect of formate concentration (s) on the specific expression rate of the three Mut phenotypes.** Strains Mut$^+$ (pSAOH5-T1), Mut$^s$ (pSAOH5-T1) and Mut$^-$ (pSAOH5-T1) were grown on minimal medium supplemented with a mixture of sorbitol (55 mM) and alanine (112 mM) with various concentrations (0-60 mM) of formate as a carbon source. Culture samples were collected in the exponential phase when the $OD_{600}$ was between 1-2, and the specific β-galactosidase activities were measured using the modified Miller assay (Materials and Methods). Specific expression rate at a particular concentration of formate was calculated by multiplying specific β-galactosidase activity with specific growth rate at that particular concentration, as shown in Eq. 3.

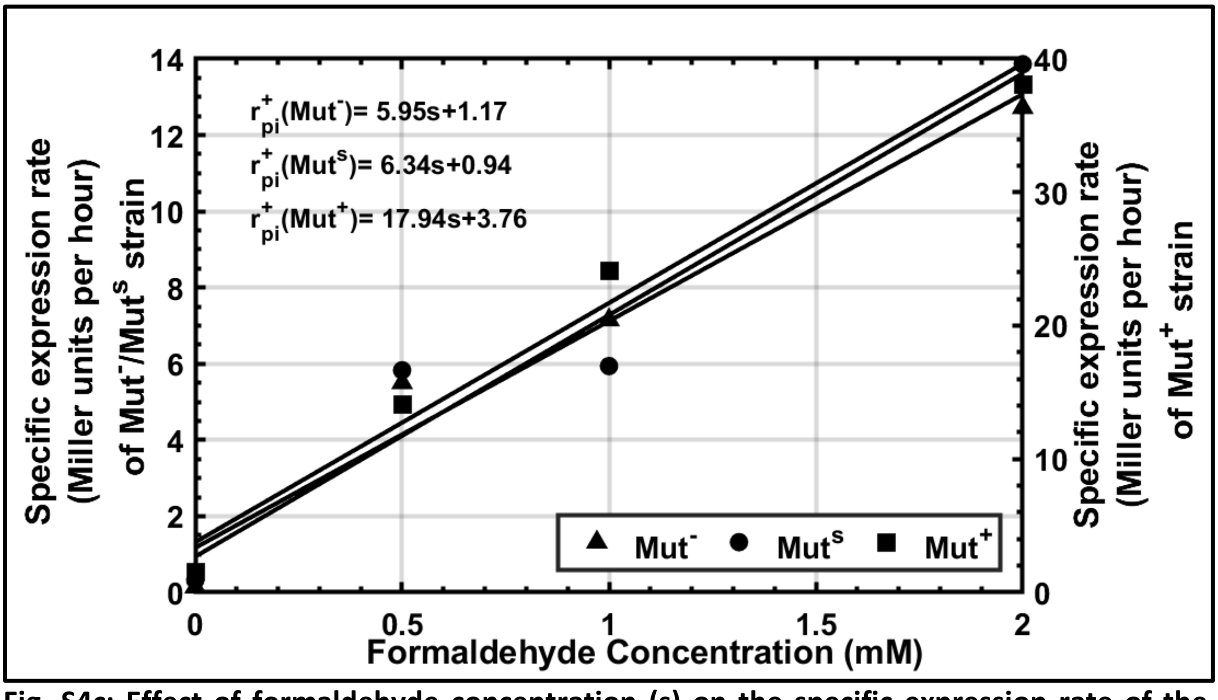


**Fig. S4c: Effect of formaldehyde concentration (s) on the specific expression rate of the three Mut phenotypes.** Strains Mut$^+$ (pSAOH5-T1), Mut$^s$ (pSAOH5-T1) and Mut$^-$ (pSAOH5-T1) were grown on minimal medium supplemented with a mixture of sorbitol (55 mM) and alanine (112 mM) with various concentrations (0-2 mM) of formaldehyde as a carbon source. Culture samples were collected in the exponential phase when the $OD_{600}$ was between 1-2, and the specific β-galactosidase activities were measured using the modified Miller assay (Materials and Methods). Specific expression rate at a particular concentration of formaldehyde was calculated by multiplying specific β-galactosidase activity with specific growth rate at that particular concentration, as shown in Eq. 3.

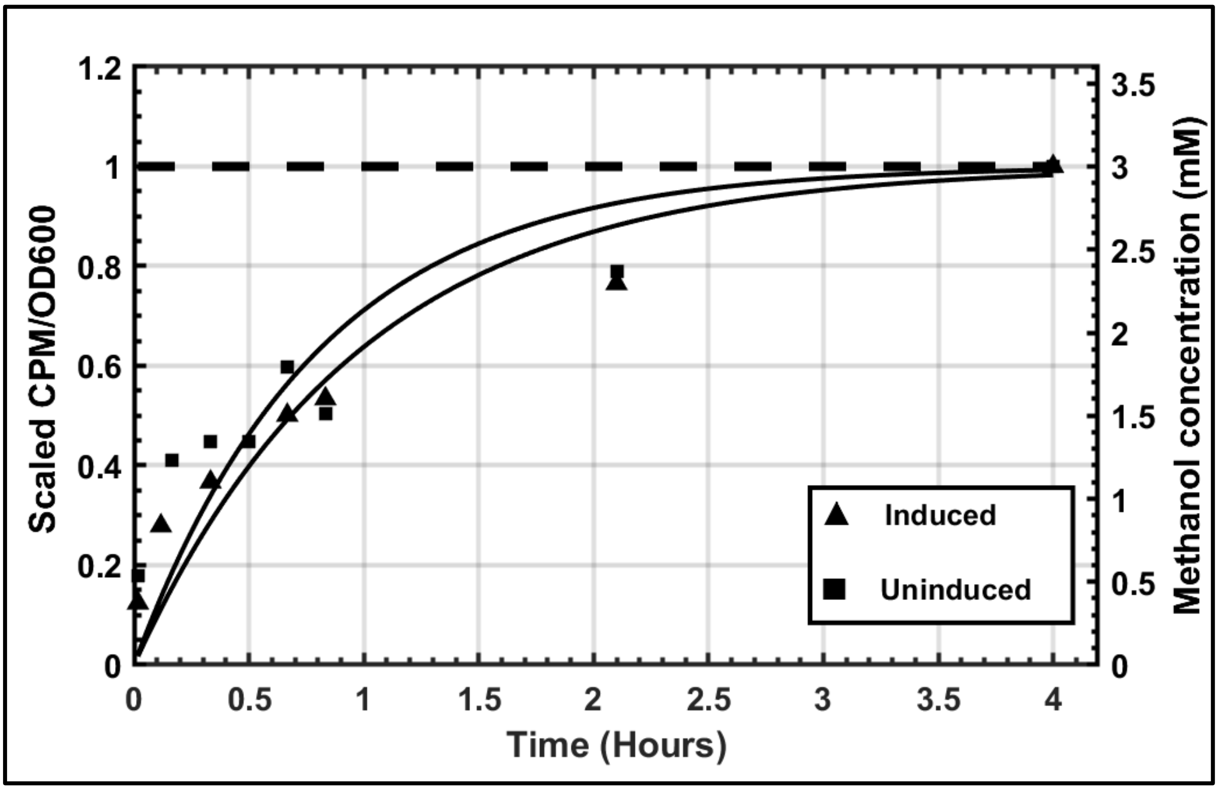


**Fig. S5: Methanol is transported by diffusion.** Strain MC100-3 was grown on either sorbitol (55 mM) + alanine (112 mM) (Uninduced) or sorbitol (55 mM) + alanine (112 mM) + formate (30 mM) (Induced) supplemented minimal medium. Cells were harvested in exponential phase ($OD_{600}$ = 0.5-1) and washed. At t = 0, radiolabelled methanol was added to both the cultures. Samples were collected at particular time intervals and immediately filtered through a membrane filter (pore size 0.45 micron). Intracellular counts were measured after washing the filters and dissolving them in a scintillation cocktail (Materials and Methods). The dotted line represents the extracellular concentration of methanol added to the medium. Intracellular concentration was calculated by using 3 x $10^{-11}$ ml as cell volume (Kunert *et al.*, 2008) and 68 % water content per cell (Kamihira *et al.*, 1987).

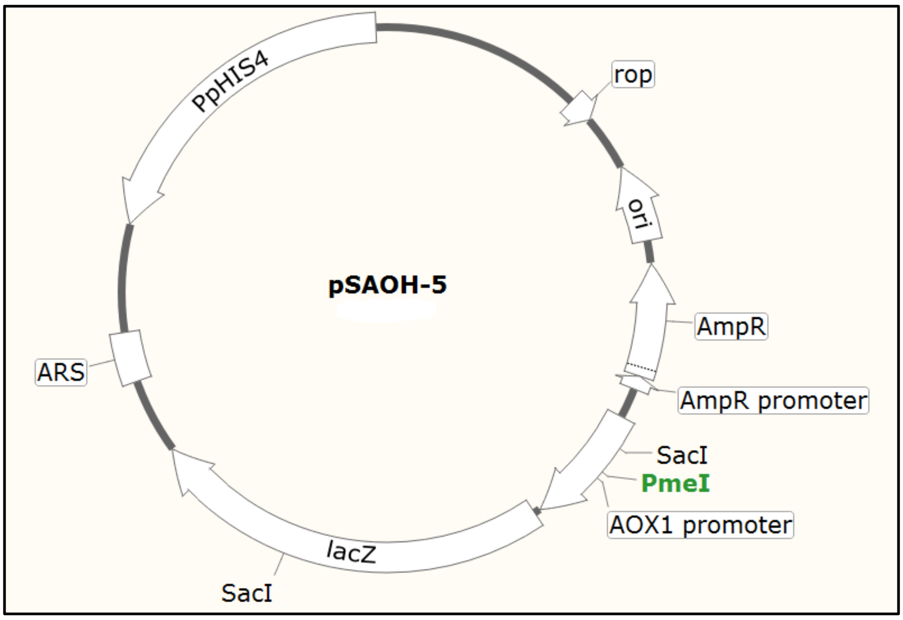


**Fig. S6: Physical map of plasmid pSAOH-5 (Tschopp *et al*., 1987a).** The unique restriction site (PmeI) in the AOX promoter sequence that was used to linearize the plasmid before transformation is indicated.

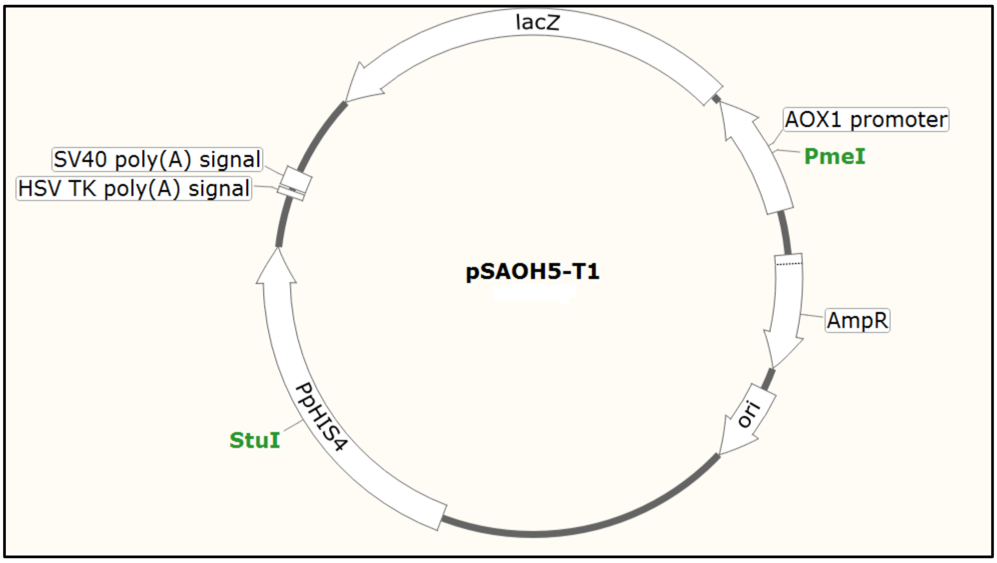


**Fig. S7: Physical map of plasmid pSAOH5-T1.** To ensure stable chromosomal integration, the ARS fragment was removed from pSAOH-5. Also, transcription terminator sequence was added at the 3’ end of the *lacZ* gene. The unique restriction site (PmeI) in the AOX promoter sequence that was used to linearize the plasmid before transformation is indicated.

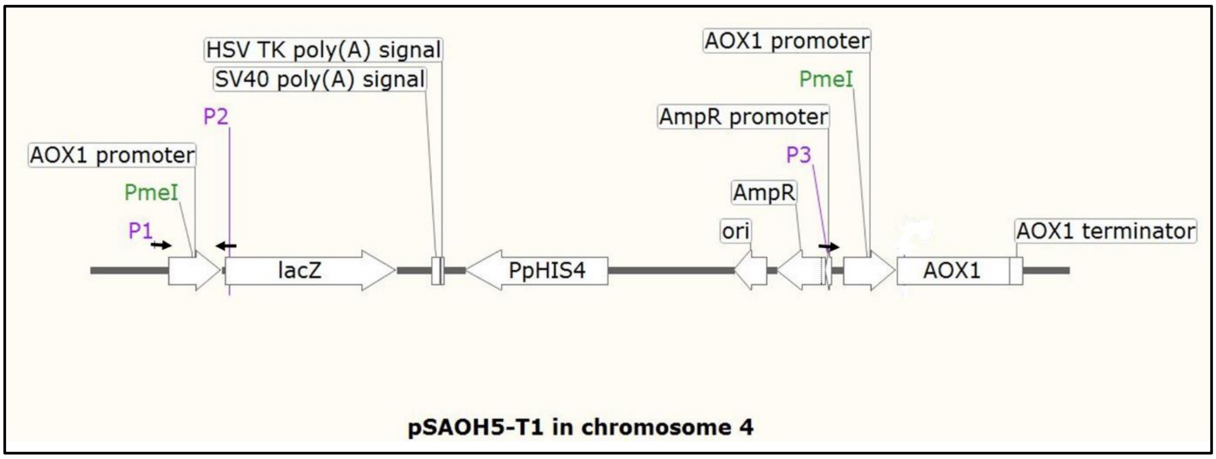


**Fig. S8: Schematic representation of a single-copy integration of pSAOH5-T1 in chromosome 4 of *K. phaffii* genome.** The plasmid is flanked by two copies of $P_{AOX1}$ at the 5' and 3' ends. Primer binding sites (P1, P2 and P3) are indicated.

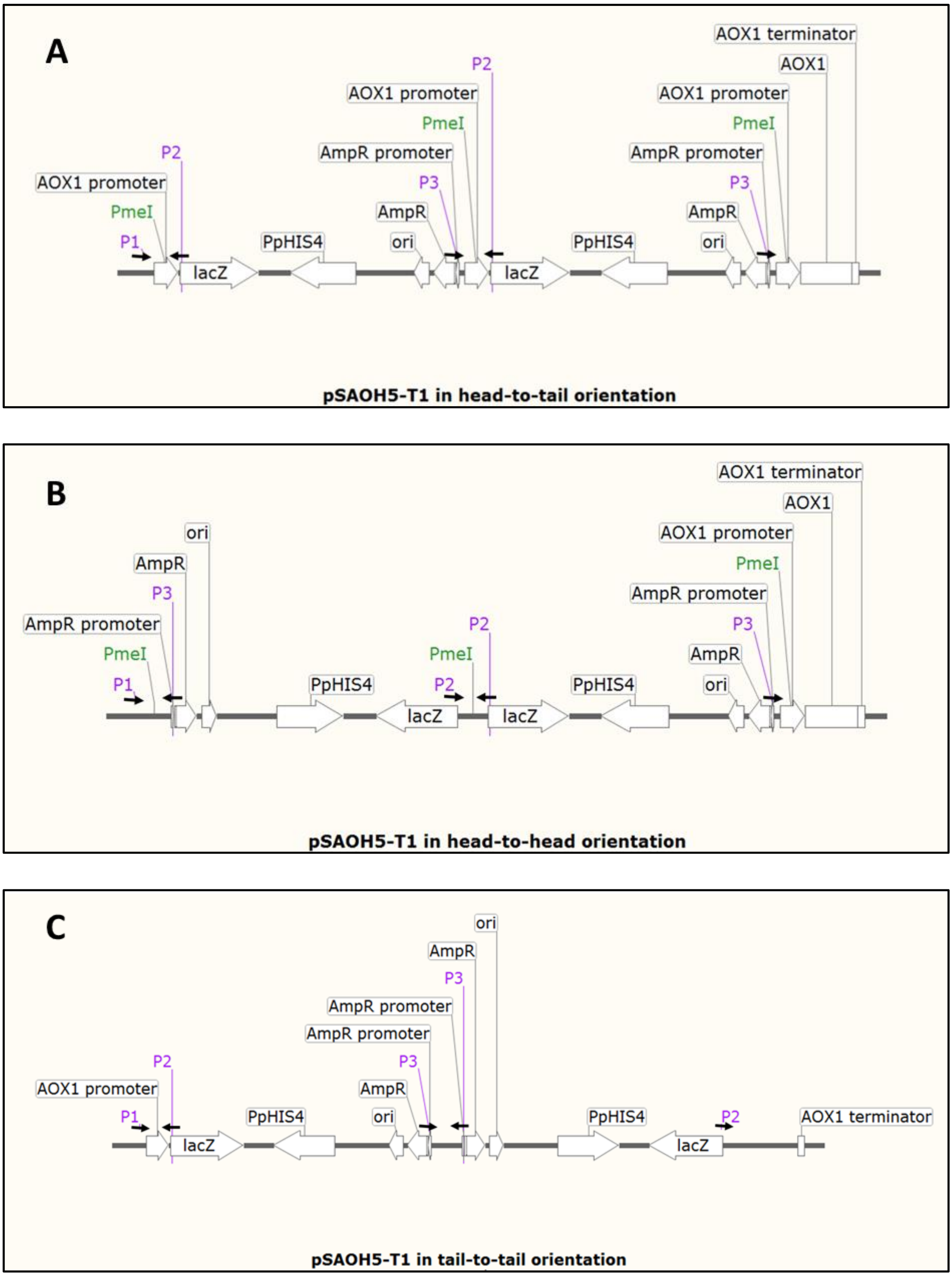


**Fig. S9: Schematic representation of multiple-copy integration of pSAOH5-T1 in chromosome 4 of *K. phaffii* genome.** Homologous recombination can lead to multiple integration events in various orientations (**A**, **B** and **C**). Primer binding sites (P1, P2 and P3) are indicated.

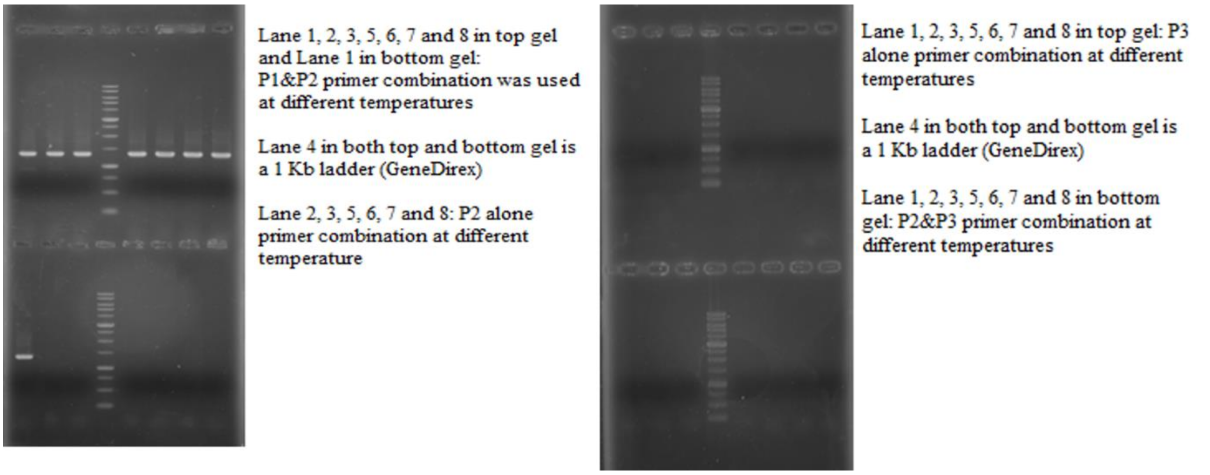


**Fig. S10: Band pattern observed with clone Mut⁻ (pSAOH5-T1) using various primer combinations.** A 1.4 kbp band was observed with primer combination P1 and P2, while no band was observed with other primer combinations (as described in Table S2). The other two strains were also confirmed in the same way (gel pictures for these not shown here).

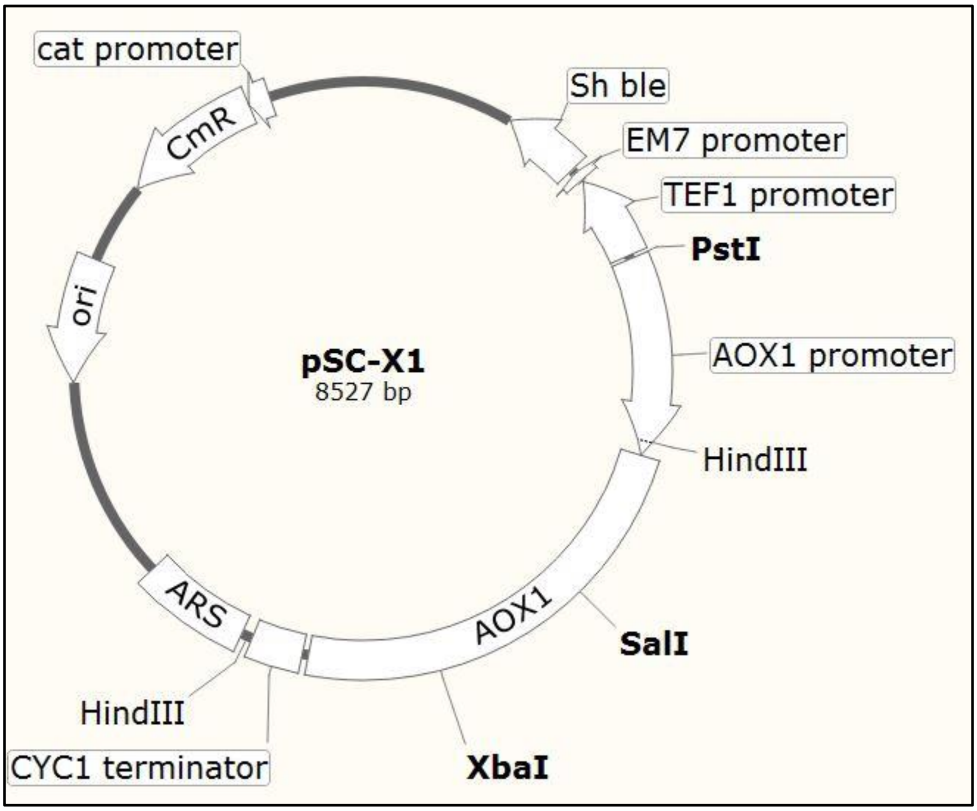


**Fig. S11: Physical map of plasmid pSC-X1.** A shuttle vector containing gene *AOX1* under its endogenous promoter $P_{AOX1}$, was constructed to express *AOX1* gene in strain Mut⁻ (pSAOH5-T1). The presence of an ARS allows autonomous replication of the plasmid in *K. phaffii* strains.

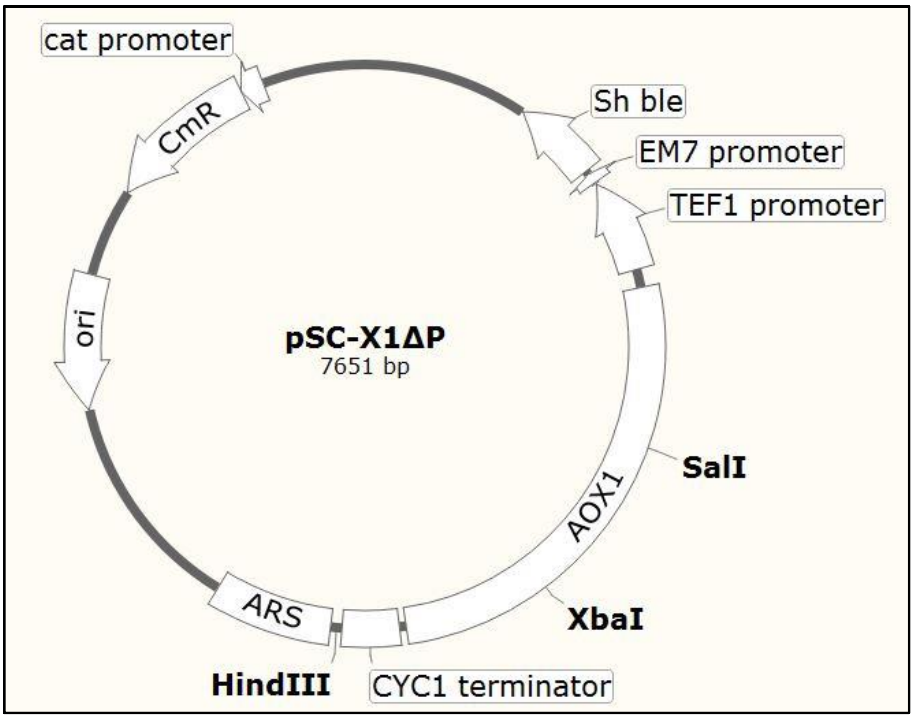


**Fig. S12: Physical map of plasmid pSC-X1ΔP.** AOX1 expression was abolished by removing the PstI-HindIII restriction fragment containing the most of the *AOX1* promoter sequence (compare with Fig. S2.6).

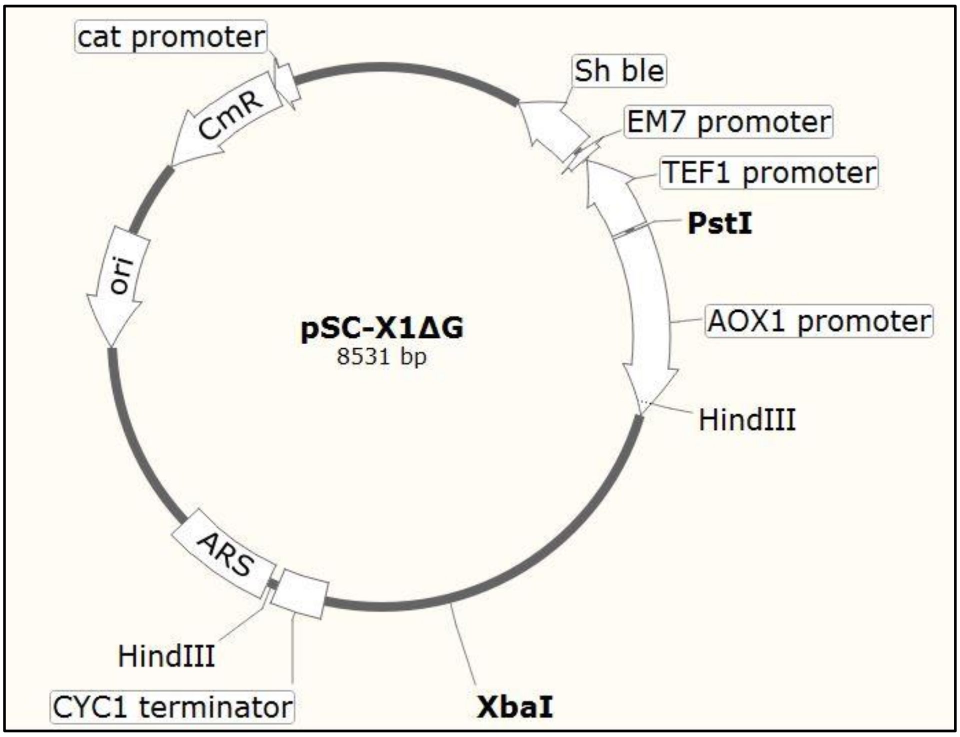


**Fig. S13: Physical map of plasmid pSC-X1ΔG.** A frame-shift mutation was introduced in the *AOX1* gene by removing the SalI restriction site from plasmid pSC-X1 (compare with Fig. S2.6)

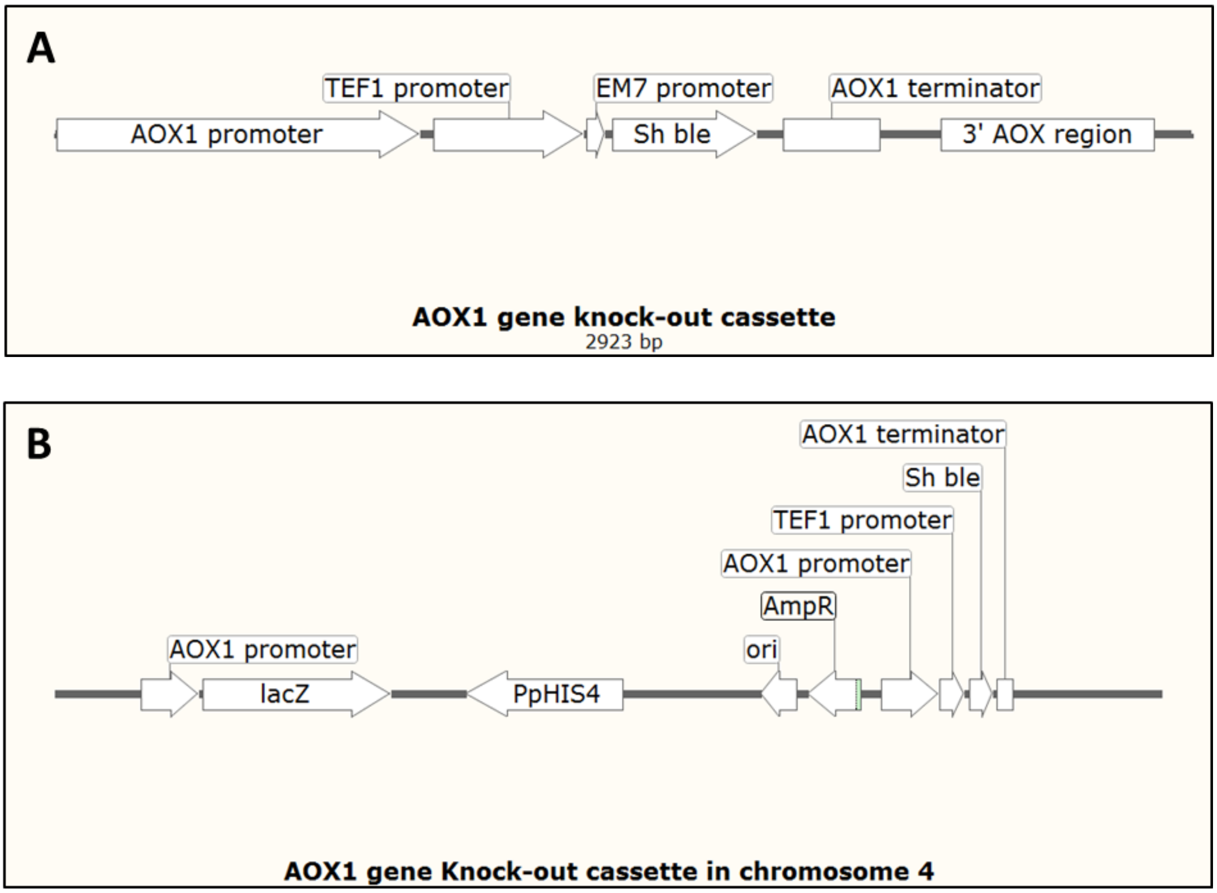


**Fig. S14: Knock-out cassette for disrupting *AOX1* gene in Mut⁺ (pSAOH5-T1). A**. Schematic representation of the knock-out cassette that was used to disrupt the *AOX1* gene of Mut+ (pSAOH5-T1). The *AOX1* gene was replaced by the *Sh ble* gene, which imparted resistance against zeocin to the transformed cells. **B**. Schematic representation of the knock-out cassette after integration into chromosome 4 at the *AOX1* locus.

#### 2. Supplementary tables

**Table S1: Specific AOX activities as measured in different Mut phenotypes and their variants.** Cultures were grown on minimal medium supplemented with sorbitol (55 mM) and alanine (112 mM) with (Induced) or without (uninduced) methanol (160 mM). Samples were collected in the exponential phase ($OD_{600}$ = 1-2) and AOX activities were measured as described in Materials and Methods.

| Induction state | $Mut^{+}$ (pSAOH5-T1) | $Mut^{-}$ (pSAOH5-T1) | $Mut^{s}$ (pSAOH5-T1) | $Mut^{-}$ (T1X1) | $Mut^{-}$ (T1X1ΔP) | $Mut^{-}$ (T1X1ΔG) | $Mut^{+}$ (T1ΔX1) |
|---|---|---|---|---|---|---|---|
| **Uninduced** | 22 ± 2 | Not detectable | 2 ± 0.35 | 34 ± 2 | 43 ± 8 | Not detectable | 2.8 ± 1.5 |
| **Induced** | 382 ± 71 | Not detectable | 4 ± 0.31 | 265 ± 20 | 36 ± 3.5 | Not detectable | 6 ± 0.8 |

**Table S2**: **PCR screening of single-integration transformants of Mut strains.** Expected DNA bands and their sizes with various primer combinations for screening for single-integration transformants of Mut strains.

| **Single Integration Event** | | |
|---|---|---|
| **Primer Set** | **No. of Bands** | **Size of Band (kbp)** |
| P1 and P2 | 1 | 1.314 |
| P2 alone | 0 | - |
| P3 alone | 0 | - |
| P2 and P3 | 0 | - |
| **Double Integration Event (Head to Tail)** | | |
| P2 and P3 | 1 | 1.343 |
| **Double Integration Event (Head to Head)** | | |
| P2 alone | 1 | 1.280 |
| **Double Integration Event (Tail to Tail )** | | |
| P3 alone | 1 | 1.406 |

**Table S3**: **Primers used in this study.**

| **Primer** | **Sequence*** |
|---|---|
| P1 | 5’-ACA TTG TCA GGA ACA CGA TG-3’ |
| P2 | 5’-TGT GCT GCA AGG CGA TTA AG-3’ |
| P3 | 5’-T ATT GTC TCA TGA GCG GAT AC-3’ |
| *AOX1*P-F | 5'-TGAC CTGCAG GATCTAACATCCAAAGACGAAAG-3' |
| *AOX1*P-R | 5'- CAA ACT CTT CGG GGA TAG CCA TCG -3' |
| *AOX1*G-F | 5'- TTCATAATTGCGACTGGTTCC -3' |
| *AOX1*G-R | 5'- GACT ACTAGT TTAGAATCTAGCAAGACCGG -3' |
| Frg.1P-R | 5'-GATC GGATCC TAG CCA TCG TTT CGA ATA ATT AG -3' |
| Frg.3-F | 5'-CTAG GATATC TCA AGA GGA TGT CAG AAT GCC -3' |
| Frg.3-R | 5'-ATCG GTCGAC ATG AGG ATC AGA CTA CGA GC-3' |

* Relevant restriction sites are underlined.
